# Programming the elongation of mammalian cell aggregates with synthetic gene circuits

**DOI:** 10.1101/2024.12.11.627621

**Authors:** Josquin Courte, Christian Chung, Naisargee Jain, Catcher Salazar, Neo Phuchane, Steffen Grosser, Calvin Lam, Leonardo Morsut

**Affiliations:** Eli and Edythe Broad CIRM Center for Regenerative Medicine and Stem Cell Research, Keck School of Medicine, University of Southern California, Los Angeles, CA, USA; Institute for Bioengineering of Catalonia (IBEC), The Barcelona Institute for Science and Technology (BIST), Barcelona, Spain; Department of Biochemistry and Molecular Biology, University of Nebraska Medical Center, Omaha, Nebraska 68198, United States; Department of Biomedical Engineering, Viterbi School of Engineering, University of Southern California, Los Angeles, California, USA

## Abstract

A key goal of synthetic morphogenesis is the identification and implementation of methods to control morphogenesis. One line of research is the use of synthetic genetic circuits guiding the self-organization of cell ensembles. This approach has led to several recent successes, including control of cellular rearrangements in 3D via control of cell-cell adhesion by user-designed artificial genetic circuits. However, the methods employed to reach such achievements can still be optimized along three lines: identification of circuits happens by hand, 3D structures are spherical, and effectors are limited to cell-cell adhesion. Here we show the identification, in a computational framework, of genetic circuits for volumetric axial elongation via control of proliferation, tissue fluidity, and cell-cell signaling. We then seek to implement this design in mammalian cell aggregates *in vitro.* We start by identifying effectors to control tissue growth and fluidity *in vitro*. We then combine these new modules to construct complete circuits that control cell behaviors of interest in space and time, resulting in measurable tissue deformation along an axis that depends on the engineered signaling modules. Finally, we contextualize *in vitro* and *in silico* implementations within a unified morphospace to suggest further elaboration of this initial family of circuits towards more robust programmed axial elongation. These results and integrated *in vitro/in silico* pipeline demonstrate a promising method for designing, screening, and implementing synthetic genetic circuits of morphogenesis, opening the way to the programming of various user-defined tissue shapes.

## Introduction

A key goal of synthetic morphogenesis in multicellular systems is to design methods to control their morphogenesis, patterning and overall functional characteristics^1–16^. One branch of this discipline aims to identify genetic circuits that could control morphogenesis by programming their self-organization. The field started in bacteria, where initial patterning circuits^17^ were followed by circuits that control proliferation, cell shape and survival to induce morphological changes in bacterial colonies ^18,19^. More recently, synthetic circuits with morphogenetic effectors have been implemented in mammalian cells^20,21^. The effectors of choice have been cadherin molecules mediating cell-cell adhesion, previously shown to guide cell rearrangements^22^. For this purpose, cadherin expression was placed downstream of synNotch receptors activation^20,21,23,24^. In another example, recombination circuits were deployed to control symmetry breaking and expression of different cadherins to achieve spheroids with different domains^25^. So far, the identification of these circuits has been done in an iterative trial-and-error approach that, although generating a plethora of different structures, does not involve forward-engineering design. Moreover, the resulting aggregates of mammalian cells in 3D have different domains but are still spherically shaped, in contrast with the elaborated architecture of real organs, which rarely display overall centrally symmetrical shapes.

We present here a combined *in vitro*/*in silico* pipeline for the design, prediction and implementation of synNotch circuits for a paradigmatic example of morphogenesis: axial elongation.

Computational approaches have proven transformational in other engineering disciplines, including bacterial transcriptional circuits^26^, leading to auto-cad frameworks for the design of complex circuits, either in hardware or software. More recently, the field has taken a step further in automation with the deployment of machine-learning and automated pipelines for complex optimization problems. We and others have developed computational frameworks to describe mammalian cell transcriptional and cell-cell communication circuits that control adhesion^25,27–30^. These frameworks have been used to predict new outcomes based on changes in initial conditions. However, so far, none of these have been used to identify, parametrize, and guide the implementation of novel circuits.

Axial elongation is recognized as one of the most fundamental operations in tissue morphogenesis, occurring during embryonic development, tissue regeneration, and metamorphosis in vivo^31,32^, and also in reconstituted stem cell-derived organoid systems^33,34,35^. It allows a spherical mass of cells to transition from radial to non-radial symmetry and is key to developing more complex structures. It is a critical milestone to achieve the programming of higher-order morphogenetic phenomena such as branching.

In the study of axial elongation in existing biological systems, since its beginning, a focus was brought to the interplay between mechanics, cell behavior, and differentiation^36^. Several specific modes of elongation have been described: convergent extension by cell intercalation^37^, tuning of cell-cell adhesion and cortical tension^38–40^, oriented cell division in drosophila wing^41^, oriented mechanical forces^42^, local control of cell proliferation^31,43–45^. In these cases, a distinction can be introduced between whether there is addition of new material (volumetric elongation) or just reorganization of existing material (non-volumetric elongation)^31^. A paradigmatic example of volumetric elongation is the anteroposterior elongation of zebrafish embryos, which seems to be supported by a fluid-to-solid, or jamming, transition^14,34,35^. Jamming transitions are ubiquitous in morphogenesis and have been found to support, among others, developmental processes^46,47^, carcinoma motility^48^ and wound healing^49^. In particular for tissue elongation, jamming transitions can provide resistance to rounding: it is in fact crucial for an elongated cell aggregate to maintain its shape as surface tension forces would promote its rounding^50,51^. Fluid-to-solid transitions can provide this resistance by increasing tissue viscoelasticity. The theoretical framework of jamming transitions explains how the properties of individual cells collectively give rise to the physical properties of tissues^52^. Cell-level properties can in turn be regulated by changes in gene expression levels or protein status. How cell properties collectively give rise to tissue physics is an active area of research. [Rigidity transitions in dev and disease - Hannezo & Heisenberg 2022]. Specific examples of cell-level properties tuning tissue-level viscoelasticity via jamming transitions are: cellular motility and actomyosin contractility^53^, the volume and composition (extracellular matrix) of intercellular space^54^, cell-matrix adhesion^32,55^, cell division and apoptosis^56^, and cell differentiation^57^. Jamming transitions in multicellular systems are complex, and effectors can have paradoxical effects; for example, increased cell-cell adhesion can either increase or decrease tissue fluidity depending on tissue cell density ^39^^39,53,58^. This inherent complexity presents a challenge for the engineer who desires to acquire effectors for specific cell behaviors to plug-and-play in their synthetic morphogenetic networks.

Synthetic gene circuits programming tissue elongation have not yet been reported to our knowledge. To address this, we asked, can we identify genetic design modules for its implementation? Furthermore, can we create complete artificial genetic circuits for elongation combining these parts? The second question underscores another compositional challenge in genetic circuit-based synthetic morphogenesis. Indeed, effectors for morphogenesis-relevant cellular behaviors have been known for a long time, and even identified as potential building blocks for more complex circuits^10,59–61^, some more than a decade ago. This somehow has not resulted in a flurry of synthetic genetic circuits for morphogenesis. At least in part, this may be due to the compositional problem: even when we have the modules that can control individual cellular behaviors, combining them in a single system is much more complicated than identifying them individually.

In this study, we first identify design rules for volumetric axial elongation through local material addition and mechanical support. We then design *in silico* a circuit that can implement that design using parameterized synthetic cell-cell signaling and effects on cell-cell adhesion, as well as changes in mechanical properties like cell proliferation and cell motility. The circuit includes a spheroid of signaling cells that activate a signal-propagation circuit in a spheroid of ‘transceiver’ cells: the activated transceiver cells reduce their proliferation and their motility, providing the mechanical support; whereas the inactivated transceivers remain proliferative and contribute the volumetric addition. Through a parameter scan screening, we identify *in silico* the most relevant parameters of this circuit to be tissue growth delta between activated and inactivated receiver cells, cell-cell adhesion, signaling speed of the signal propagation, and - to a certain extent - tissue fluidity between activated and inactivated cells. We then identify and characterize, *in vitro*, effectors genes to support elongation via repurposing available parts (synNotch signal propagation circuit^62^, cadherin expression for cell sorting through adhesion control^21^), and via screening of candidates for control tissue growth (e.g. p21) and tissue fluidity (e.g. CA-RhoA and CA-MLCK). We then move to combine these modules *in vitro* in complete synthetic elongation circuits following two main strategies: distinguished by either constitutive or inducible N-cadherin expression in the transceivers. Finally, we parametrize our *in silico* model with the novel effectors and generate a morphospace where we localize both the parametrized *in silico* simulations, and the implemented *in vitro* circuits; this helps us identify potential implementation augmentation for future improvements of this first circuit, in particular along the lines of identification of effectors (or other external boundary conditions) to counteract endogenous tendencies of these cell spheroids to round. We think that this work represents a further step towards the rational design of morphogenetic circuits in multicellular mammalian systems.

## Results

### *In silico* design of a gene circuit for axial elongation

Natural systems are an inexhaustible source of inspiration for engineers. To extract design principles to guide the implementation of volumetric elongation we looked at relevant examples across several domains. In these examples, elongation is obtained through the local, distal addition onto a rigid “stalk”, of a non-rigid material; the non-rigid material is then transformed into a further segment of the stalk itself by a rigidification process. Material addition and rigidification proceeds in parallel (**Fig. 1a-b**). Examples from which we extracted these designs include: the elongating zebrafish presomitic mesoderm, where cells migrating from the dorsal medial zone enter a polarized location of the distal end of the presomitic mesoderm (gray), which stiffens (red) in a gradient towards the embryonic anterior pole^53^. In the growth of a stalagmite, dissolved calcite is deposited by water drops at the top of the structure following a gravity gradient (gray), calcite which then precipitates to form the rigid body of the stalagmite (red)^63^. In bone elongation, cells in the growth plate divide and self-renew providing polarized addition of material (gray), before differentiating in osteoblast which forms the bone structure (red)^64^. This design has been implemented also in a man-made robot, the FiloBot^65^, where asymmetrically deposits of molten polymers implement volume addition (gray), and their progressive hardening (red) provides a structure on which more molten polymer can be deposited, thus leading to elongation. The common theme of those very distinct examples is that material is locally and in a polarized manner added in a fluid-like region (grey), before converting (white arrows) into a more rigid matter (red regions), which provides structural support for the local deposition of more material. The polarized conversion of the new material to more rigid constitutes the basis for directional elongation.

**Figure 1.**
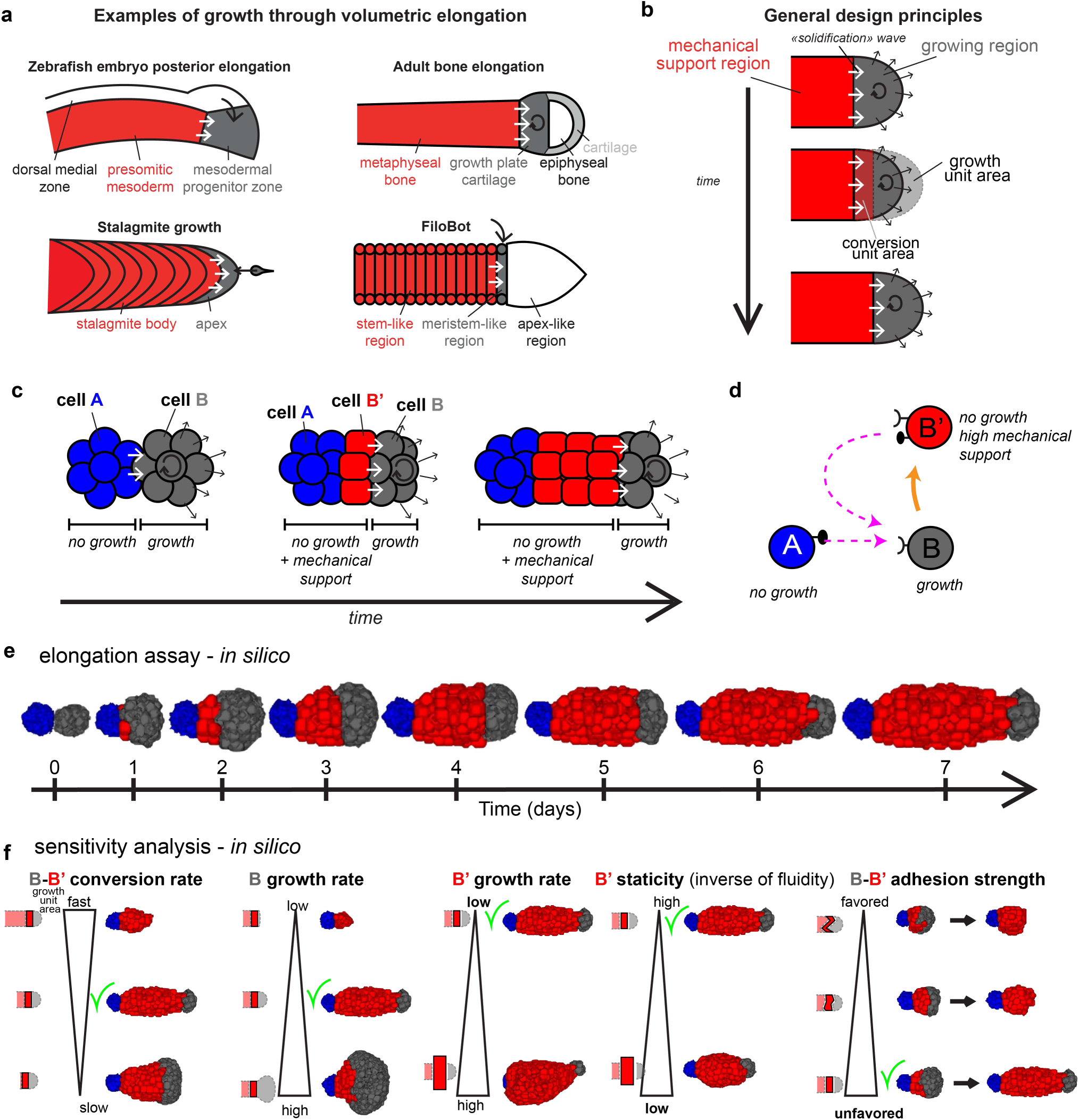
Volumetric elongation logic and computational implementation. **a.** Four schematized examples of volumetric elongation: top left, zebrafish embryo posterior elongation; top-right, bone elongation; bottom left, stalagmite growth and bottom right, Filobot - see Main text for details and references. For all the examples, red is the mechanically rigid structure; gray is the more fluid region where there is an addition of mass (black arrow); and the extension of the rigid structure with the incorporation of a part of the gray added mass is indicated with white arrows. **b.** Schematic depicting the underlying design principles of a type of volumetric elongation. Shown are snapshots over time (vertical arrow) of an elongating structure; the gray region is expanding in the direction of the black arrows at the same time as the red region is expanding towards the region of the gray mass that is most proximal to it. Highlighted in the second snapshot are the growth region per unit of time, and the conversion region per unit of time. When the 2 are equal, sustained elongation can proceed. **c.** Schematic representation of an implementation with cellular units of a rule-based circuit that could implement volumetric elongation. Cell type A (blue) and cell type B (gray) are initially in two juxtaposed spheroids; cell type B’ (red) emerges at the interface between A- and B-type cells. **d.** Schematic representation of the rules followed by the cells in c. **e.** Snapshots taken every 24h of simulated time over 7 days of a computational implementation of the rules shown in d., realized in the GJSM CC3D framework. **f.** Selected results from the sensitivity analysis (the complete analysis is in Fig. S1). On top of each triangle is noted the varied parameter. On the left of the triangles is a schematic depiction of how the growth speed of the proliferative B area (gray area), the growth speed of the rigid B’-type cell compartment (red), the B->B’ conversion speed (light red) and the shape of the B-B’ interface (red-gray interface) change as the parameter in study changes. On the right of the triangles are the day 7 snapshots of how the structures derived with the corresponding parameter variation look like. Green ‘check’ signs are noted close to the examples of successful elongation.

Starting from this high-level description of rules for this specific implementation of volumetric axial elongation, we wanted to generate a realization in a multicellular morphogenetic circuits design environment we recently developed: GJSM (General Juxtacrine Signaling Model)^27^. Briefly, the computational system is based on a Cellular Potts system, further parameterized with baseline proliferation, timing, adhesion and synNotch signaling parameters obtained from *in vitro* data obtained with engineered L929 cells. This allows us to: (i) generate ‘realistic’ simulations, and (ii) suggest direct genetic circuits to implement in cells. In this environment, we can simulate cell-cell juxtacrine signaling, cell-cell adhesion, cell state changes, cell proliferation, and cell motility. This framework was developed to easily translate *in silico* circuits into *in vitro* implementations relying on the synNotch receptor^27^.

To identify potential synNotch-based circuits, we derived cell-based rules from the design principles outlined above (**Fig. 1c**). The initial conditions consist of a spheroid of rigid, non-proliferating cells (A, blue) in contact with a spheroid of fluid and proliferating cells (B, gray). Cells B proliferate to provide material addition, and also convert to activated cells (B’, red) upon direct contact with the A-type cells, or with already activated B’-type cells. B’ type cells are less proliferative and collectively form a structurally rigid structure. Thus, the B-B’ conversion wave should result in a rigidification wave, whereas the remaining B-type cells should provide the needed continuous material addition.

To implement this type of conversion of B-type cells to B’-type, we designed the synNotch-compatible circuit logic of **Fig. 1d**: A-type cells signal to B-type cells to activate to a B’ state; and the B’ state is also capable of activating more B-type cells to the B’ state. In this way, the signaling wave could be supported by an available synNotch signal propagation circuit^66^. From this description, we generated an initial circuit in the GJSM system, given it was already parametrized for cell-cell synNotch signaling and cell-cell adhesion. For proliferation and cell motility, which were previously parametrized only for parental L929 cells, we used extreme values of no motility and no proliferation for B’-type cells (for complete *in silico* genome, see genome #1, **Table 1.1**). As shown in **Fig. 1e**, this implementation resulted in a cellular system that displays sustained elongation over the course of 7 simulation days, leading to the formation of a growing, narrow elongating structure: at the beginning, the surface of contact between the 2 spheroids is an area of signaling from the A-type cells to the B-type cells, that turn into B’-type cells only at the interface. As gray B-type cells keep proliferating, they also keep turning into red B’-type cells due to the activation signal coming from older B’-type red cells to their left.

Overall, these *in silico* simulations show that a circuit relying on juxtacrine signal propagation and cell state change from a proliferative state to a non-proliferative and collectively rigid state could achieve axial elongation from the initial conditions of fusing two spheroids of different identities.

### *In silico* sensitivity analysis identifies modules for *in vitro* implementation of axial elongation

We sought then to use this successfully implemented *in silico* system of axial elongation to identify which modules were the most important for *in silico* elongation. We performed a sensitivity analysis for the following parameters (**Fig. S1**): adhesion strength between A, B, and B’-type cells (6 pairs: 6 parameters), proliferation and motility in each cell type (6 parameters), B-B’ conversion rate (1 parameter). Each of the parameters at play was switched from one extreme to another (for example, completely stopping the proliferation of the B-type cells, or having the B’-type cells proliferate as much as the B-type cells), to have a qualitative understanding of which parameters are more impactful to the phenotype (genomes #2-#19 in **Table 1.1**, also indicated in **Fig. S1**).

As shown in Fig. S1, some parameter modifications barely affect the elongating phenotype, such as increasing the motility of A-type cells or increasing the strength of A-B’ adhesion (**Fig. S1b**). Another set of parameter changes gives rise to mild changes; increased B’ motility, reduction of B->B’ conversion speed, and increase of B or B’-type cell proliferation (**Fig. S1c**). A third group of parameter changes generate initially elongated structures that though exhaust the gray cap of proliferating B-type cells, so the elongation ceases after a certain initial phase of elongation: weaker A-A adhesion, increased B->B’ conversion speed, and reduced B’-B’ adhesion (**Fig. S1d**). Finally, a group of perturbations prevented any discernible elongation from occurring: reduced B-type cell motility, reduced B-type cell proliferation, weaker B-B adhesion, stronger B-B’ adhesion and stronger A-B adhesion (**Fig. S1e**).

Focusing on the cell of type B-B’, the parameters that mostly impacted phenotypes are (**Fig. 1f**): B->B’ conversion rate, ratio between B and B’ growth rate, B’ motility and B-B’ adhesion strength.

From these results, we aimed to identify design principles to guide the implementation.

1. We found it important to balance the conversion and growth (**Fig.1b**). This can be affected by altering B->B’ conversion rate, and/or the volumetric growth of B/B’-type cells: if the rate of conversion of B-type cells to B’-type cells is too fast, or if the B-type cells do not replicate fast enough, the conversion area is bigger than the growth area, and the proliferating mass gets consumed before any elongation can be achieved. Conversely, if the conversion rate is too slow, the conversion area is smaller than the growth area, and the proliferating region overgrows, leading to an elongating but widening structure.
2. Another important design principle is to control the shape of the B-B’ conversion zone, which needs to remain relatively smooth and flat to in turn instruct the polarization of the newly added material. We found that to reach this goal we should maintain a low heterotypic B-B’ adhesion compared to homotypic B-B or B’-B’ adhesion. This results in a neat separation between the B-type cells’ “cap” and the B’-type cells “stalk”. Increasing B-B’ adhesion results in an increasingly rugged and convex B-B’ interface, overall increasing B-B’ contact area and thus conversion rate through juxtacrine signaling (**Fig. S1f**).
3. Another design principle is to have a high rigidity of the B’-type cell region. Indeed, increasing B’ motility (which should increase tissue fluidity^38^), and weakening B’-B’ adhesion (**Fig. S1c-d**), have modest but noticeable detrimental effects on elongation. This is in line with previous results and theory, as tissue viscosity (opposite of fluidity) resists surface tension which rounds up non-spherical tissues over time^53^.

Overall, these *in silico* analyses alongside the studies on natural elongation systems cited in the introduction, suggest that important modules for tissue elongation include controlled signal propagation, controlled cell proliferation, controlled tissue rigidity, and controlled cell-cell adhesion.

### *In vitro* design

The circuit we designed in the GJSM computational framework was planned so the same circuit logic could be implemented *in vitro* with synNotch-based modules. As shown in **Fig. 2a**, A-type cells would express a synNotch ligand (e.g. GFP-lig). The B-type cells express the cognate anti-GFP synNotch receptor, which activates several modules upon stimulation (conversion to B’-type cells): GFP-ligand itself, for encoding the signal propagation module; and effector genes to control cell adhesion, cell proliferation, and tissue fluidity (**Fig. 2b**). With the appropriate effectors, once spheroids of A-type cells and B-type cells are formed, at the interface A-type cells should communicate with the first neighbors on B-type cells in the B-type cells spheroid and turn them into B’-type cells characterized by GFP-lig expression and activation of effector genes; these B’-type cells then will activate their B-type cells neighbors in a polarized manner, on the right in the figure (**Fig. 2c**).

**Figure 2.**
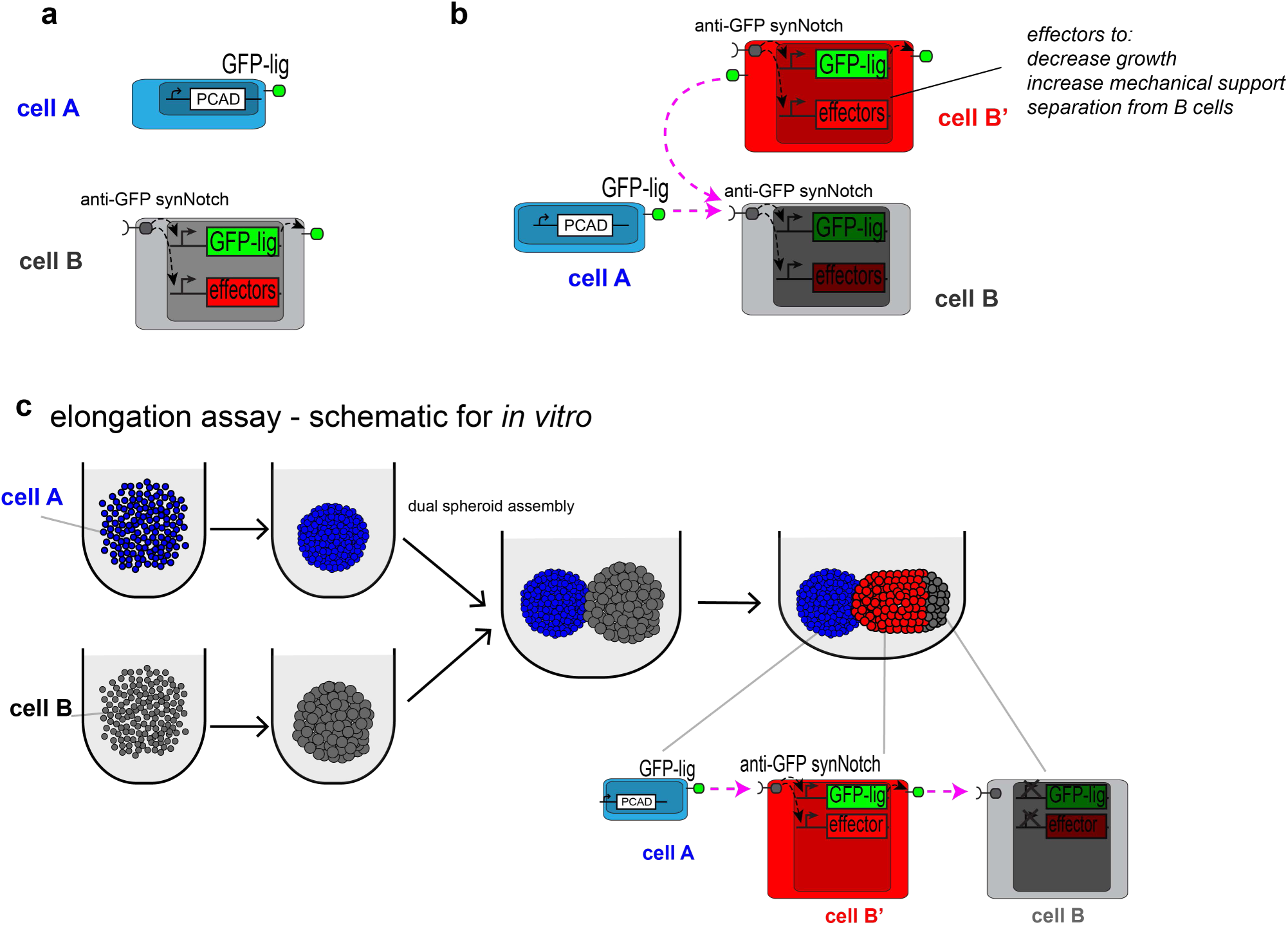
Design of a topology of genetic circuits for the implementation of volumetric elongation. **a.** Schematic depiction of the logic of the circuit in cells. Cell A and cell B, with their artificial genetic parts. Cell A contains an adhesion molecule (P-cad) and GFP-lig; cell B contains an anti-GFP-synNotch receptor that activates GFP-lig (green) and cell biological effectors (red). **b.** Schematic of the activation of cell B to B’ by cell A. Dashed purple arrows indicate activatory ligand/receptor interactions. Specific genetic effectors are discussed in the following sections of the paper. **c.** Schematic setup for an “elongation assay”. Cells A (blue) are dispensed in a U-bottom well in media (light gray) and left to aggregate for 24h, after which they appear as a compacted blue ball. Similarly for gray B-type cells. At this point an assembloid of the 2 spheroids is created by pipetting both spheroids in the same U-bottom well. Given the opportunity to interact, the A-type cells would activate the B-type cells to turn into B’-type cells (red) at the interface between the blue spheroid of cell A, and the gray spheroid of cell B. At the bottom are indicated the cells and the signaling relationships.

We noted from the start that, although it would be ideal to consider the four modules (signaling, adhesion, proliferation and fluidity) as orthogonal to each other, the reality of cellular and tissue biology is far from a perfect set of orthogonal components. To a certain extent, all modules interact with each other, but it is especially when it comes to tissue fluidity that the effectors are most complex to identify. So, we first identify separated modules for signaling, adhesion and proliferation; for the tissue fluidity, we consider it in combination with the others. In the following paragraphs we describe how we characterize the available modules for signaling and adhesion, we identify novel effectors for proliferations, and then for tissue fluidity; for each module, we first describe an assay that we used to score the different circuits or effectors, and then how we picked each of those.

### Signaling module

We first focused on the signaling module. The signaling module is a signal propagation circuit where “transceiver” (B/B’-type) cells in the inactive state (B-type) are able to be activated by neighboring cells presenting the signal (e.g. cell type A), upon which they activate and become B’-type cells, themselves able to activate their neighbors. In the *in silico* implementation, this part of the circuit allows the signal to be propagated from the sender A cells through the spheroid of B cells in a wave-like fashion.

We have previously implemented this circuit in L929 cells and studied its patterning capacities in 2D^62^. Briefly, transceiver cells are generated by lentiviral-mediated integration of constructs to express anti-GFP/tTA synNotch receptor under a constitutive SFFV promoter, and tTA-responsive promoter driving transcription of GFP-lig upon receptor activation; clonal populations are then derived from the initially polyclonal infected population. In order to obtain a range of signal speeds, we derived clonal populations from the initially polyclonal infected population of these transceiver cells. The sender (A-type) cells, which constitutively express GFP-lig and the nuclear far-red protein H2B-miRFP703, were maintained as a polyclonal population after infection. When triggered by sender cells, a transceiver cells ensemble can propagate the signal activation across a field of cells in 2D cultures^66^. To test if this circuit would work in 3D, we generated 2 spheroids of 1000 cells each, one of sender (A-type) cells which expressed GFP-lig (alongside a far red reporter and the adhesion molecule P-cadherin for increased compaction, see more on compaction below); and one of transceiver (B-type) cells. These two spheroids are then assembled in a single ultra-low attachment U-bottom well plate, and the signaling is allowed to occur over a time course of several days (**Fig. 3a-b**). As shown in **Fig. 3c**, the transceiver spheroid is activated upon contact with the sender spheroid: GFPlig expression increases first at the point of contact, before spreading to the rest of the spheroid. We repeated the same setup with multiple transceiver clones. In **Fig. 3c-d**, we report the result from a “slow” clone and “fast” clone: the slow clone has a signal propagation rate of roughly 0.6um/h, whereas the fast clone has a propagation rate of 2um/h (**Fig. 3d**, see also **Fig. S2e** for single channel view of the images). As a control, B-type cells activate only minimally by day 2 by themselves (**Fig. S2b**). We also show that when propagation assays are done in presence of increasing doses of doxycycline (which inhibits the synNotch-mediated gene expression) this results in a decrease of signal propagation speed, as expected (**Fig. S2a**). Of note, in all these experiments, the overall structure of the two-spheroids assembloid proceeds to become a sphere after an initial phase where the 2 individual spheroids are still discernible in a dumbbell shape.

**Figure 3.**
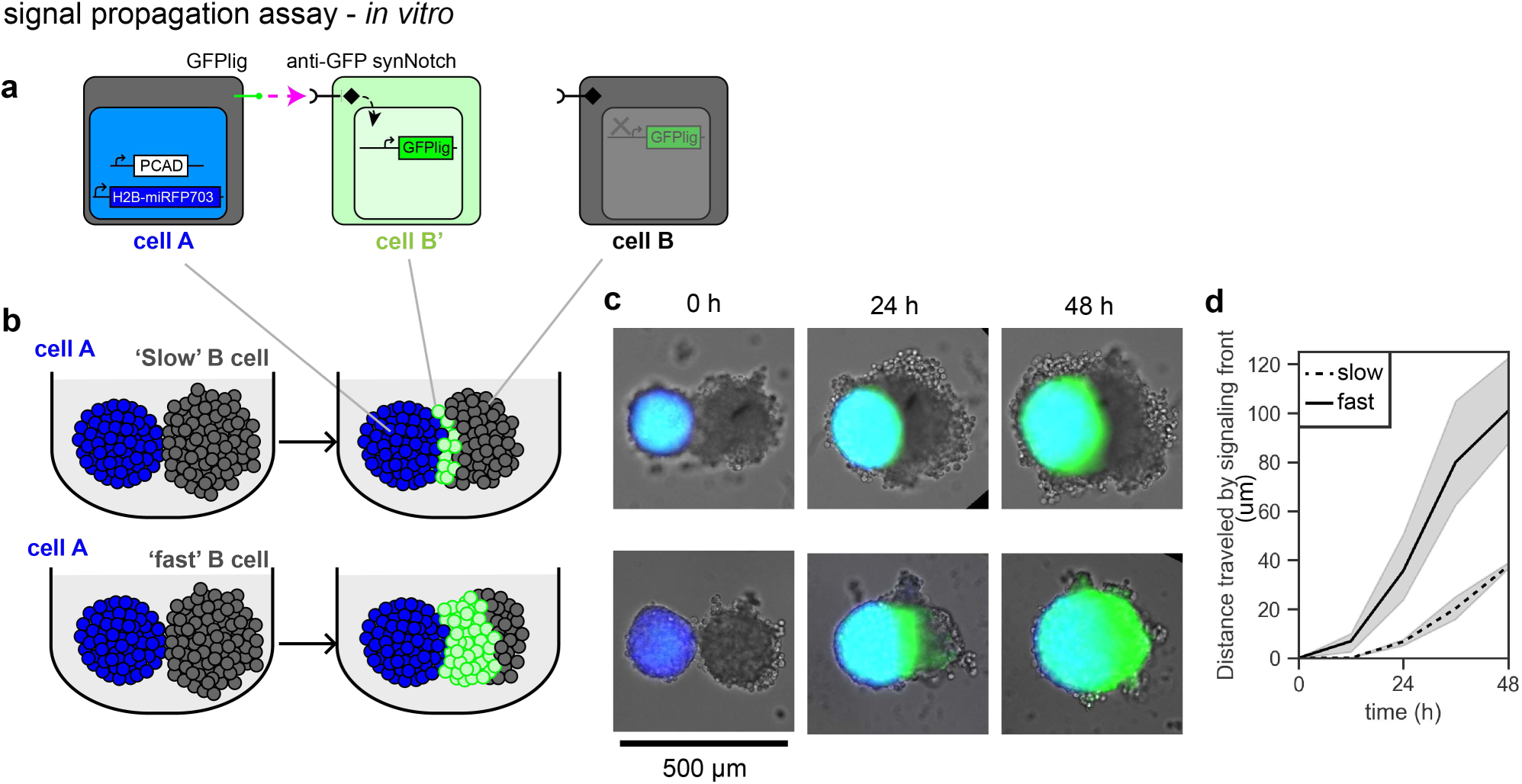
Implementation *in vitro* of a synNotch based circuit for signal propagation in 3D. **a.** Schematic depiction of cells of type A, B, and B’. H2B-miRFP703 is a nuclear far-red fluorescent protein. In this case, B-type cells only activate GFP ligand in response to activation, no effector genes. **b.** Schematic depiction of a signal propagation assay in 3D for a slow-propagating clone (top) and a fast-propagating clone of B-type cells (bottom). **c.** Fluorescent microscope images at the indicated time points during a signaling propagation assay, using the cells from the schematic in a; at the beginning, 1000 cells of type A or B are used to generate the spheroids. The bright-field image is overlaid with the blue and green signals; the far red fluorescent protein signal is rendered in blue; the GFP ligand is rendered in green. See videos SV5-6. A-type cells are a polyclonal population; each B-type “clone” is from a clonal population. **d.** Graph of distance traveled by signal over 48h of propagation assay for the slow clone (dashed line) and the fast clone (continuous black line). For the method of calculation of the front from the imaging dataset see Methods. (n=3-4 technical replicates).

Collectively, these results show that we have a module for signal propagation that works in a 3D environment with the initial condition of two adjacent spheroids, and that can be modulated, as required for the circuit to sustain axial elongation.

### Adhesion modules

Cell-cell adhesion mediated by cadherins is shown to affect cell positioning in L929 cells, both by simple overexpression^22,67^ and by synthetic circuit expression^21,25^. Moreover, it may affect spheroid compaction, proliferation, fluidity and signaling. In fact, in the *in silico* circuit for spheroid elongation presented in **Fig. 1**, changes in adhesion in the B cells are critical for sustained elongation.

We thus performed a series of experiments to test some the effects on L929 spheroids and on the signaling module first of changes in cell-cell adhesion. A first requirement for our system is that the initial condition would bring together two rather compact spheroids, that do not intermix with each-other, so that the signal propagation is initiated from a clean and localized interface between A-type and B-type cells. To achieve that, we tested if two known cadherins (N-cad and P-cad) could support this organization, by lentivirally overexpressing the cadherins with constitutive promoters in L929 cells and forming 1000 cells spheroids from the sorted polyclonal populations. As shown in **Fig. 4a**, both P-cad and N-cad overexpression led to better compaction of spheroids following initial seeding, achieving one of the design specifications. To test what this configuration would result in morphology-wise when two such spheroids are juxtaposed, we generated a assembloids from 2-day old spheroids made with 1000 N-cad+ cells and spheroids made with 1000 P-cad+/miRFP+ cells, and evaluated their morphological reorganization over 6 days. The fact that P-cad+ cells have also a fluorescent marker allowed us to follow cellular reorganization alongside global morphological changes. To quantify variability among different assembloids, we quantified the global aspect ratio (AR) of the co aggregate: AR is calculated as the ratio of the major axis over the minor axis, and is a value that scores 1 for circles, and >1 for more elongated ellipses (See Fig. S7 for methods of AR quantification). As seen in **Fig. 4b-c**, assembloids made with same-cadherin spheroids (i.e. N-cad + N-cad or P-cad + P-cad) resulted in the formation of an ellipsoidal structure in 1 day, and of a spherical structure in 3 days, reaching AR values close to 1, characteristic of spherical structures. The heterotypic assembloid where P-cad+ spheroid is juxtaposed to N-cad+ spheroid resulted instead in a stabilization of a structure with an aspect ratio around 1.3 throughout day 6. Of note, in all cases the first day the AR is elevated is due to the experimental initial conditions that have the two spheroids just barely touching (**Fig. S2c**).

**Figure 4.**
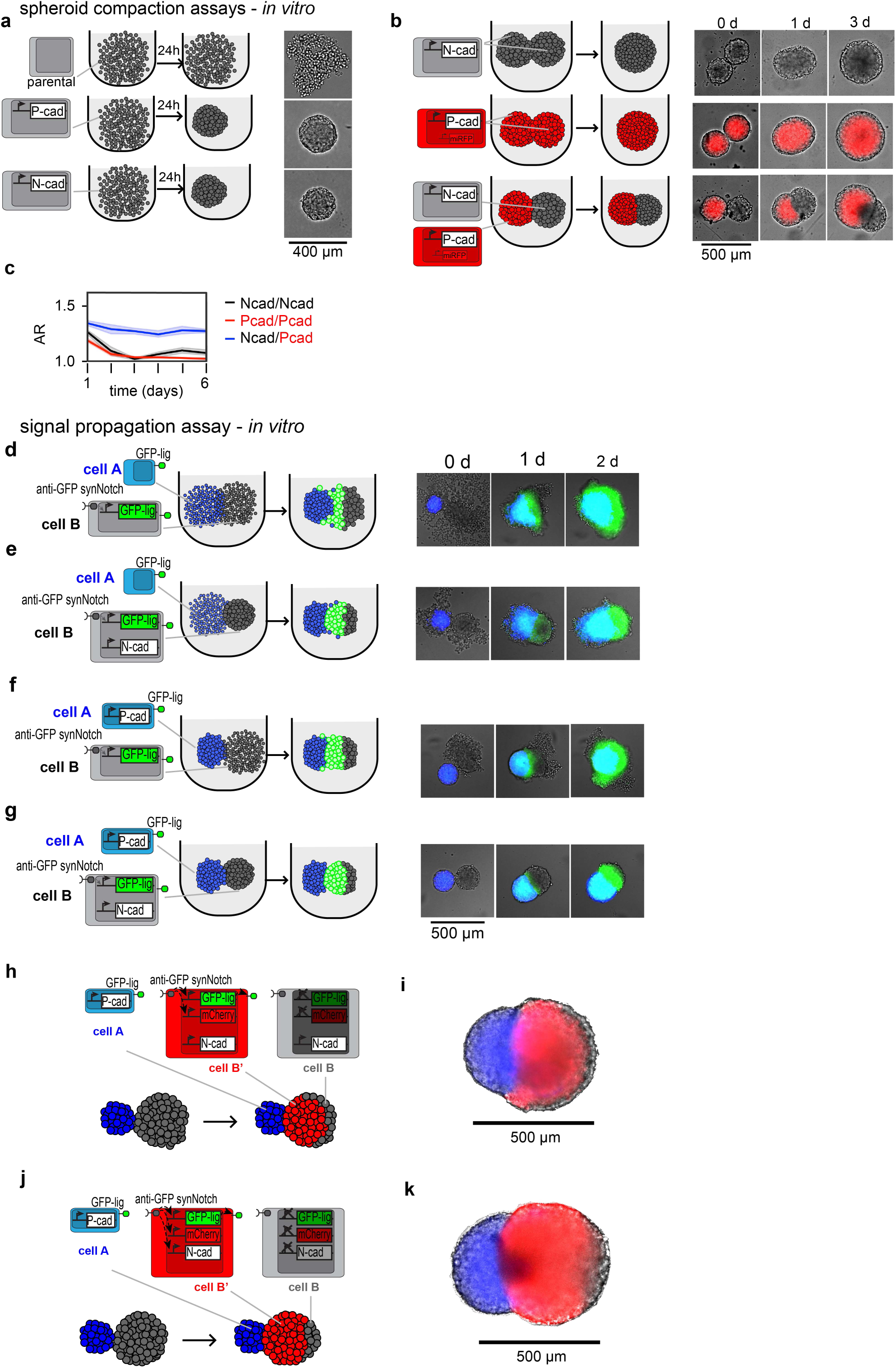
Expression of adhesion proteins in cells A and B affects the compaction of the aggregates and the signaling propagation. **a.** Schematic (left) and microscope bright field images (right) for a spheroid compaction assay. On the far left is a schematic of the cell used in the assay in the 3 different rows: top, parental L929 cells; middle, L929 overexpressing P-cad; bottom, L929 overexpressing N-cad. 1000 cells are aggregated on a U-bottom microwell per each condition and left to aggregate for 24h. Scale bar 500um. **b.** Schematic (left) and microscope bright field and fluorescent images (right) for an assembloid compaction assay. On the far left is a schematic of the cell used in the assay in the 3 different rows: top, gray cells, N-cad spheroid + N-cad spheroid; middle, red cells, P-cad spheroid + P-cad spheroid; bottom, red cells + gray cells, P-cad + N-cad spheroid. On the right, time-stamped microscope images of the corresponding experiments; gray cells are overexpressing N-cad; red cells are overexpressing P-cad; the red signal is from miRFP703 (miRFP) expressed in P-cad+ cells. Scale bar 500µm. **c.** Graph showing quantification of the aspect ratio (AR) of the assembloid compaction assay corresponding to the experiments illustrated in b. For details on methods of quantification see Methods and Fig. S7. **d-g.** Signal propagation assay as in Fig. 3a-b; left, schematic depiction of the cells, the initial condition and the observed propagation; right, microscope images taken at the indicated time (0d = day 0, …) after assembloid of 1000 A-type cells and 1000 B-type cells are left to aggregate for 48h. The far-red signal in the A-type cells is rendered in blue; the green signal from activated B’-type cells is rendered in green. In **d.**, A-type and B-type cells do not overexpress any adhesion protein; in **e.** B-type cells overexpress N-ca d; in **f.** A-type cells overexpress P-cad; in **g.** A-type cells overexpress P-cad and B-type cells overexpress N-cad, as indicated in the schematics. See also Supplemental Videos SV1-4. **h-i.** Consequences of the constitutive overexpression of adhesion proteins for the propagation front. **h.** Schematic of a signal propagation assay with A-type cells (blue, starting with 200 cells) overexpressing P-cad, and B-type cells (gray, starting with 4,000 cells) overexpressing N-cad, and inducing GFP-lig, and mCherry reporter in the B’ state (red). **i.** Segmented microscope image with overlaid bright field, red channel (mCherry in B’-type cells), and blue channel (far red protein in A-type cells); B-type cells are the gray cells on the right that are not red. Non-segmented micrographs and individual channels are depicted in Fig. S3. **j-k.** Consequences of the inducible overexpression of adhesion proteins for the propagation front. **j.** schematic of a signal propagation assay with A-type cells (blue, starting with 200 cells) overexpressing P-cad, and B-type cells (gray, starting with 4,000 cells) inducing GFP-lig, and mCherry reporter, and N-cadherin only in the B’ state (red). **k.** Segmented microscope image with overlaid bright field, red channel (mCherry in B’-type cells), and blue channel (far red protein in A-type cells); B-type cells are the gray cells on the right that are not red. Non-segmented micrographs and individual channels are depicted in Fig. S3.

We then asked if and how cadherin expression in both spheroids would affect signaling. To do so, we compared results obtained with a sender (A-type) cell line constitutively expressing or not expressing P-cad, and a fast transceiver (B-type) cell line constitutively expressing or not expressing N-cad. These cell lines were generated started from sender cells and fast transceiver cells upon which lentiviral integration of P-cadherin or N-cadherin lentiviral particles respectively, and sorting cadherin expression. With these cells, we generated spheroids of 1000 cells and aggregated them pairwise; the results are shown in **Fig. 4d-g** (see also **Fig. S3a-d**). When the assembloids are generated from spheroids without overexpressed cadherins, pipetting the transceiver spheroid to bring it in contact with the sender spheroid results in a loss of its integrity, with many loose cells peripheral to the assembloid. The signaling activation starts as the two spheroids intermix extensively, and by day 2 all the transceiver cells are activated (**Fig. 4d**). Similarly, when either only A-type cells or only B-type cells overexpress a cadherin, all the B-type cells are converted to B’-type cells in 2 days, and we still observe loose cadherin-negative cells attaching to various regions of the assembloid (**Fig. 4e,f**). On the contrary, when A-type cells have P-cadherin, and B-type have N-cadherin, the propagation front is the most linear, and the two spheroids have a more compacted and aggregated structure, with no disaggregated cells (**Fig. 4g**, see also **Fig. S2d** for controls); moreover, some B-type cells remain inactivated at day 2, on the most distal end of the transceiver spheroid.

*In silico* experiments suggested that phase separation between B’-type and B-type cells was important for constraining tissue growth to the distal end of the system and maintaining a stable signaling interface between transceivers, and that changes in the adhesion matrix would affect the shape of the propagation interface (**Fig. S1f**). We checked if expression of constitutive or inducible adhesion molecules in cell B would affect propagation interface; as shown in **Fig. 4h-i** (and **Fig. S3e**), constitutive N-cad in B-type cells in a propagation assay with 200 A-type cells and 4,000 B-type cells results, at day 6, in a propagation front that is convex, with inactivated B-type cells distributed along the surface of the spheroid, distal to the A-type cells spheroid. Instead, when we built B-type cells where anti-GFP synNotch induces N-cadherin alongside GFP-lig, a flatter propagation front where the inactivated B-type cells are constrained at the right pole **Fig. 4j-k** (and **Fig. S3f**). For the generation of the N-cadherin inducible B’-type cells, lentiviral integration of a construct for TRE→N-cad was used that renders N-cadherin inducible downstream of synNotch activation. This phenotype was shown by *in silico* experiments to be driven by low B-B’ adhesion compared to B-B or B’-B’ adhesion (**Fig. 1e**) and thus suggests that inducible N-cad in transceivers can lead to some level of B-B’ segregation compared to its constitutive expression.

Collectively, these results show that expression of cadherin affects spheroid compaction, assembloid geometry and signal propagation, and that the most linear signal propagation geometry is obtained with A-type cell spheroid with P-cadherin and B-type cell spheroid with N-cadherin, either inducible or constitutively expressed.

### Cell proliferation module

We have identified modules for signaling, adhesion and aggregation following the specifications informed by the computational model predictions to obtain axial elongation. To do so, we have repurposed parts (synNotch circuit and cadherins) that were already available and partially characterized in L929 cells. For the other two remaining modules, proliferation and tissue fluidity control, we did not have off-the-shelves effectors. We thus set up assays and small-scale screens of candidate effectors to identify them.

For proliferation, we devised an *in vitro* assay to screen genetic effectors that can achieve changes in the volumetric growth rate of 3D tissues (See Methods and **Fig. S4**). We reasoned that the re-implementation of genetic parts in different cellular context could be problematic, so we decided to perform the screening in the basic chassis of the transceiver (B-type) cells constitutively expressing N-cad (except when N-cad expression was itself the tested effector), i.e. an anti-GFP synNotch expressing cell that produces GFP in response to activation under a TRE promoter. We engineered this basic cell line with TRE promoters regulating a small library of effectors that are reported to decrease proliferation when overexpressed (**Fig. S4a.i**); these polyclonal cell lines, derived from the clonal fast transceiver line, are passaged in the presence of tTA inhibitor tetracycline to prevent any activation of the effectors outside of the experimental setup. To activate effector overexpression and test their effect on cell proliferation, we plated cells on plate-adsorbed GFP ligand, which resulted in activation of the transgenes (both GFP-lig and the effector genes, see **Fig. S4a.ii**). We then generated spheroids and assessed the volumetric growth of the spheroids over time and compared it with the volumetric growth of the same cell line cultured with doxycycline (from 2 days before seeding), which abolishes synNotch signaling and effector gene expression. To do so, we measured spheroid diameter over 5 days, inferred the volume of the spheroid, assuming its spherical shape, obtaining a growth index metric reflecting the amount of daily volumetric growth of the spheroid: 0 for a spheroid with no growth at all, 1 for a spheroid whose volume doubles every day (see methods for detailed equations).

We identified and cloned 4 different cyclin-dependent kinase inhibitors (CDKIs), p21, p27, p16, and p53^68–70^. We engineered B-type cells constitutively expressing N-cad to activate, in the induced B’ state, mCherry (as a control) or the candidate CDKI effectors (and mCherry). We also engineered B-type cell lines to activate N-cad, or N-cad+p21 when induced to the B’ state. We then generated spheroids made of each of those cell lines in the activated (B’-type) or non-activated (B-type) state and measured their growth index (**Fig. 5a-b**). As shown in **Fig. 5 c**, we found that while basic B-type cells with no or mCherry control effectors have a similar growth rate when activated or not, inducing the 4 CDKIs significantly inhibited volumetric growth when the cells were turned in their B’-type state. The extent of growth inhibition differed between the 4 effectors, p21 reduced the growth index from 0.35 to 0.26, p16 to 0.27, p27 to 0.19, and p53 to 0.12. The strongest growth index reduction in a single experiment, with freshly generated ip53 transceivers, was down to 0.05 (**Fig. 5a-b**). N-cad, on the other hand, was found to increase the growth index from 0.31 to 0.39, whereas simultaneous N-cad + p21 induction reduced growth in the cell line (**Fig. 5c** and **Fig. S6d**). Across the board, we observed that effector efficiency decreased with successive passages (not shown). Importantly, mCherry induction levels were in the same range among effectors, suggesting that their differential impact on cell proliferation was not strictly due to a difference in inserted copy number (**Fig S5a,c**).

**Figure 5.**
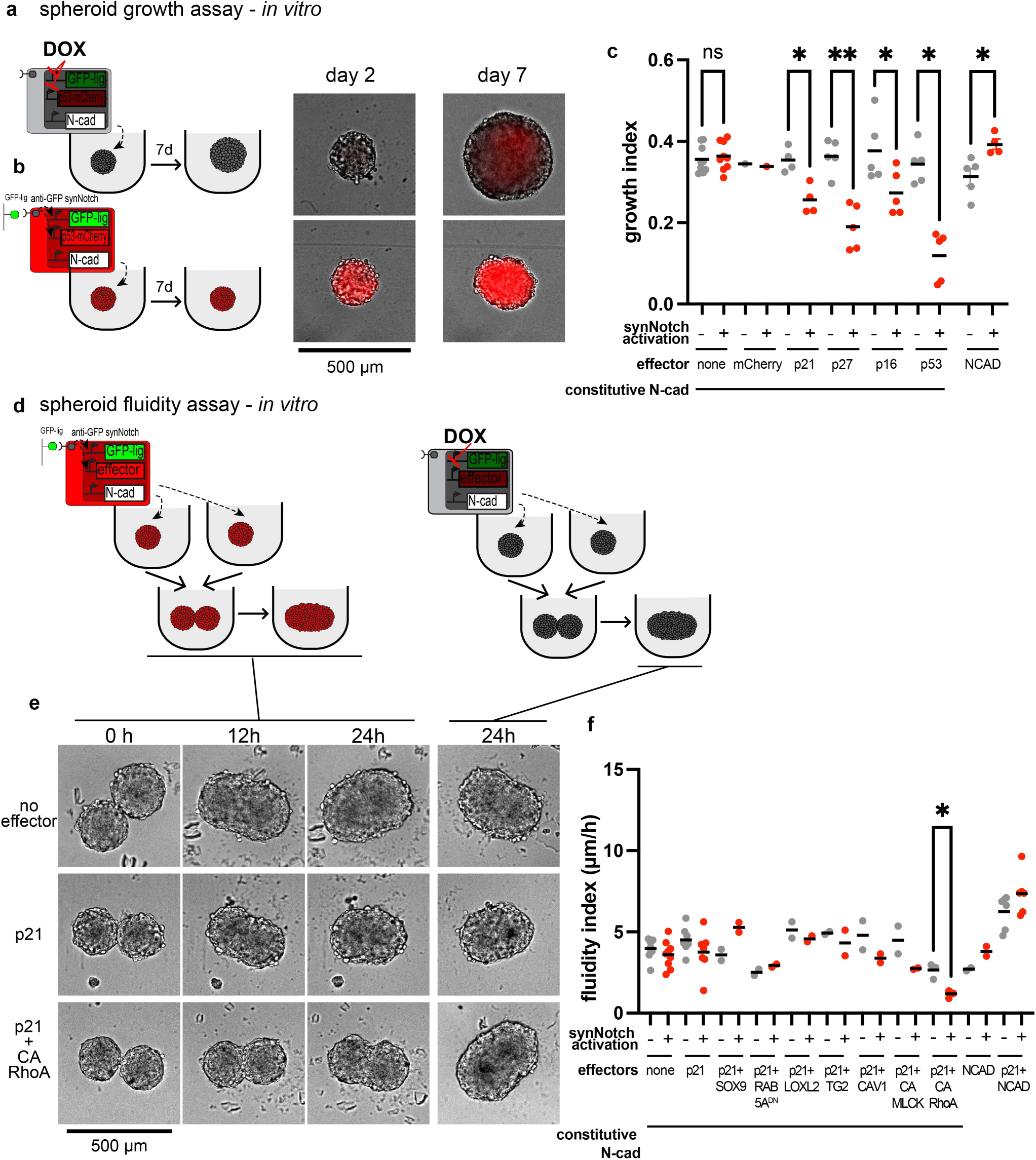
Identification of synNotch pathways for inhibition of spheroid growth and tissue fluidity. **a-b.** Spheroid growth assay: 1000 transceiver (B-type) cells are dispensed in a U-bottom well, and their growth is captured by imaging on days 2 and 7. On the right, representative microscope images of a spheroid of B-type cells with inducible p53, either preactivated (B’-type) by culture on GFP, or inhibited (B-type) by dox at the indicated time points. mCherry signal is rendered as red. Scale bar 500um. **c.** Growth index graph for the indicated conditions. Per each effector, a condition without synNotch activation (dox, gray dots), and one with synNotch activation (via plate-bound GFP, red dots) are reported. For details on methods of growth index quantification see Methods. Growth index of 0 means no growth at all; growth index of 1 means duplication of volume every day. **d.** Schematic of spheroid fluidity assay. For more details, see Fig. S3. Briefly, transceiver (B-type) cells with diverse inducible effectors were preactivated or preinhibited before 1000-cell spheroid seeding as explained above. The resulting spheroids were brought together in homotypic pairs and imaged every hour for 24 hours. **e.** Microscope images of spheroid fluidity assay with no effectors (top row), inducible p21 (middle row), and inducible p21+CA-RhoA (bottom row). The first three columns (0h, 12h, 24h) are brightfield images taken from activated (B’-type) cells. The fourth column (24h) represents brightfield images taken from inactivated B-type cells of the same genetic effectors. Scale bar 500um. **f.** Fluidity index graph for the indicated conditions. Per each effector, a condition without synNotch activation (dox, gray dots), and one with synNotch activation (via plate-bound GFP, red dots) are reported. For details on methods of fluidity index quantification see Methods and Fig. S3d. High fluidity index means the spheroids are fluid and tend to fuse fast; low fluidity index means the spheroids are more viscous and tend to fuse more slowly. For **c.** and **f.** Brown-Forsythe and Welch ANOVA test results: * is p<0.1; ** is p<0.01 all other differences between measurements in ON vs. OFF transceivers were non-significant.

We also measured sender A-type cells in this volumetric growth assay and calculated that the growth index of senders is roughly twice as high as that of transceivers, with a DGR of 0.68 ± 0.05 (n=3 ± std.dev.).

Thus, we identified and characterized pathways that, in response to synNotch ligand GFP, affect L929 spheroid volumetric growth.

### Tissue fluidity module

We then moved to identify and characterize genetic effectors of tissue fluidity. To evaluate tissue fluidity, we used a previously reported^50^ spheroid fusion assay, where the speed at which two spheroids made from the same cells fuse, depends on the viscocapillary speed (or viscocapillary velocity) of the system (in µm/h) (**Fig. 5d** and **Fig. S4c-d**). This metric is increased by surface tension and opposed by tissue viscosity^50,71^, hence we call it here the ‘fluidity index’, which is inversely proportional to the viscosity of the system: a lower fluidity index means that the system will keep a non-spherical shape longer, which is what we aimed for.

Given that proliferation could affect fluidity also, we first performed fluidity assays with the proliferation effectors discussed earlier. As shown in **Fig. S6a**, induction of p16, p27, and p53 exhibited a trend of increased tissue fluidity or unchanged fluidity, whereas overexpression of p21 tended to decrease fluidity to almost half of the fluidity of ON ip53 spheroids. We then moved to combine a well-performing proliferation effector (p21) with a screen of effectors that, based on a literature review, would impact the properties of the cytoskeleton or membrane or ECM to increase tissue viscosity. The effectors we chose include: the SRY-box transcription factor 9 (SOX9), a master transcription factor for chondrogenesis^72^; caveolin 1 (CAV1), a promoter of cell-level rigidity^73^; a constitutively active mutant of myosin light chain kinase (CA-MLCK^74^) and a constitutively active mutant of the Rho GTPase RhoA (CA-RhoA^75^), both major regulators of actomyosin contractility, impacting cell contractility and motility^76^. We engineered cNCAD fast transceivers to overexpress those genes (in combination with p21) when activated (**Fig. 5d**). To achieve this design, as previously, we co-infected L929 transceiver cells with lentiviral constructs for TRE>p21 alone, or TRE>p21 + TRE>fluidity effector (see more in methods) and derive polyclonal cell lines expressing the transgenes. We then performed fluidity assays with spheroids made with these cell lines. As shown in **Fig. 5e-f**, we found that there was a light trend for naive transceivers and p21 transceivers to decrease fluidity when activated, a trend for p21+SOX9 to increase tissue fluidity, a trend for p21+CAV1 and p21+CA-MLCK to decrease tissue fluidity, and a statistically significant decrease in fluidity when p21+CA-RhoA were co-induced. When we tested induction of N-cad, either alone or in combination with p21, we instead observed a trend for increased fluidity index upon activation (**Fig. S6c**).

To complete the characterization of these cell lines with a combination of p21 and tissue fluidity effectors, we measured the effect of effectors activation on tissue growth and found that tissue growth was reduced in all these transceiver lines, consistent with the effect of p21 seen when activated alone (**Fig. S6b-d**).

Thus, we identified and characterized synNotch pathways that, in response to ligand GFP, concurrently decreased proliferation and decreased tissue fluidity. Based on the computational simulations this would mean mechanical support by the B’-type cells in the elongation circuit.

### *In vitro* implementation, complete circuits, ‘cytoskeletal strategy’

As we generated a control circuit for compacted spheroid reduction of tissue growth and fluidity upon signaling transceivers activation, we decided to evaluate the phenotypic outcomes of complete circuits, where the signaling, adhesion, proliferation, and tissue fluidity modules are assembled.

We divided our attempts into two categories. In the first class of circuits (**Fig. 6**), sender A-type cells constitutively express P-cad, and B-type cells fast clone express constitutively N-cad and, in response to synNotch activation, they activate GFP-lig in addition to either p21 alone or p21 in combination with the fluidity effectors CA-MLCK or CA-RhoA. The second class of circuits, presented in the next section (**Fig. 7**), instead relies on induced N-cad expression.

**Figure 6.**
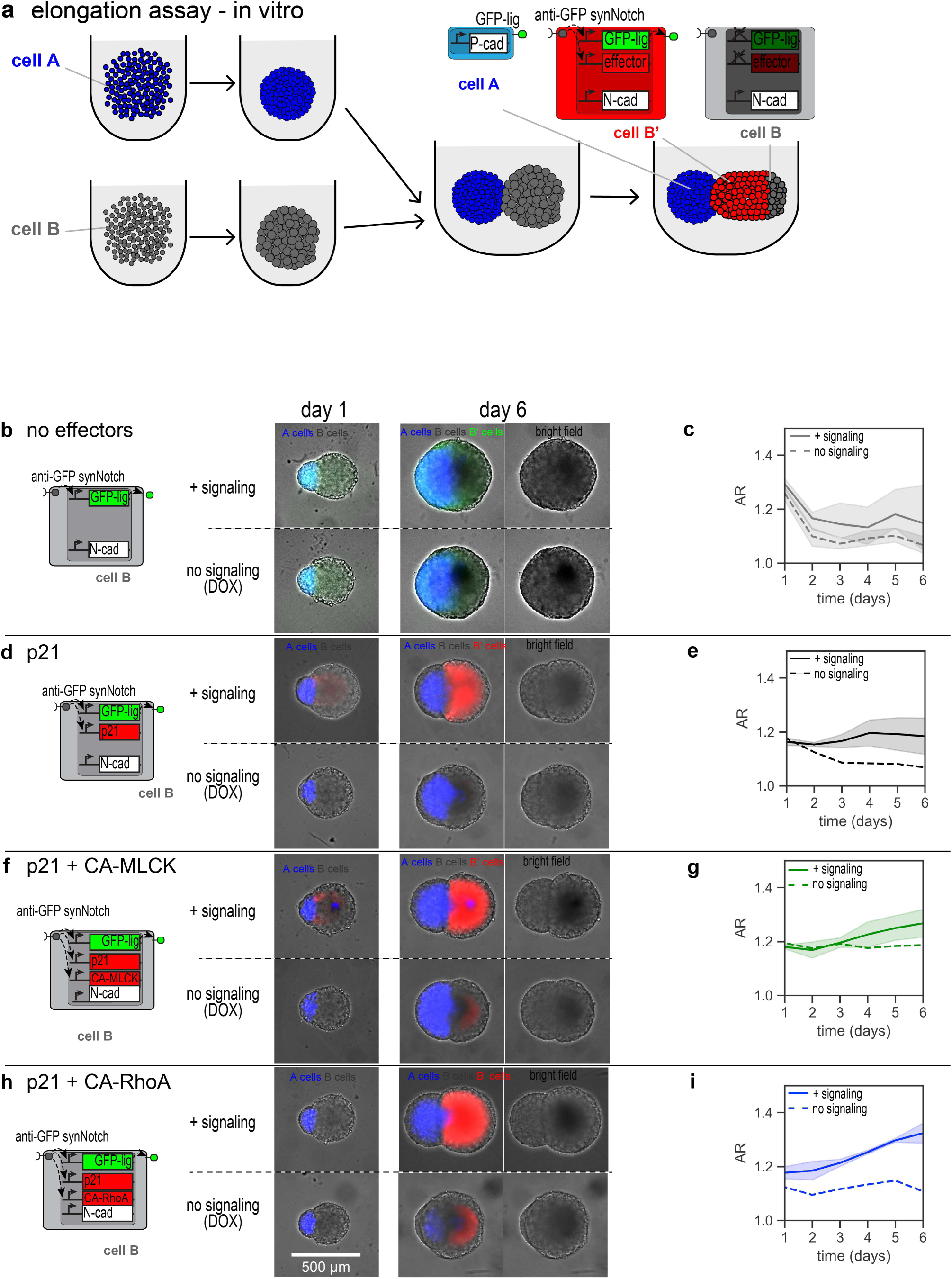
Programming tissue elongation through dynamical control of tissue fluidity and growth. **a.** Schematic of elongation assay: spheroids of 200 sender (A-type) cells (blue) and 4,000 transceiver (B-type) cells (gray) are seeded separately. All B-type cells in this figure have constitutive N-cadherin, anti-GFP-synNotch that activate GFP-lig for signal propagation, and one or more inducible effectors as indicated below. After 2 days, the two resulting spheroids are assembled in an individual well, and the assembloid is imaged daily henceforth. **b.** Elongation assay with B-type cells that activate only mCherry reporter and no effectors. Left, schematic of B cell circuit. Right, micrographs of Day0 and Day6 of two conditions: top row, condition where synNotch signaling in B-type cells is allowed to happen; bottom row, condition where synNotch signaling in B-type cells is impaired with the small molecule Dox. In the micrographs the far-red signal from the constitutive far-red marker in A-type cells is rendered in blue, and the GFP signal is rendered in green. Scale bar is 500um (shown at the bottom, valid for all panels). **c.** Graph of aspect ratio (AR) over time for n=1 experiment, n=3-6 technical replicates. The continuous black line is for samples with signaling; the dashed black line is for samples with DOX, i.e. without signaling. AR =1 for a circle, AR>1 proportionally to the elongation of the ellipsoid. For more details on AR calculation see methods. **d.** Elongation assay with B cells that activate p21 as proliferation effector. Left, schematic of B cell circuit. Right, micrographs of Day0 and Day6 of two conditions: top row, condition where synNotch signaling in B cells is allowed to happen; bottom row, condition where synNotch signaling in B cells is impaired with the small molecule Dox. In the micrographs the far-red signal from the constitutive far-red marker in A cells is rendered in blue, the mCherry signal in B’ cells is rendered in red, and the GFP signal is not shown. Scale bar is 500um (shown at the bottom, valid for all panels). **e.** Graph of aspect ratio (AR) over time for n=2 +signaling experiments and n=1 no signaling experiment, each averaged from 3-6 technical replicates. The continuous black line is for samples with signaling; the dashed black line is for samples with DOX, i.e. without signaling. AR =1 for a circle, AR>1 proportionally to the elongation of the ellipsoid. For more details on AR calculation see methods. **f.** Elongation assay with B cells that activate p21 and CA-MLCK upon synNotch signaling activation. Left, schematic of B cell circuit. Right, micrographs of Day0 and Day6 of two conditions: top row, condition where synNotch signaling in B cells is allowed to happen; bottom row, condition where synNotch signaling in B cells is impaired with the small molecule Dox. In the micrographs the far-red signal from the constitutive far-red marker in A cells is rendered in blue, the mCherry signal in B’ cells is rendered in red, and the GFP signal is not shown. Scale bar is 500um (shown at the bottom, valid for all panels). **g.** Graph of aspect ratio (AR) over time for n=2 +signaling experiments and n=1 no signaling experiment, each averaged from 3-6 technical replicates. AR =1 for a circle, AR>1 proportionally to the elongation of the ellipsoid. The continuous green line is for samples with signaling; the dashed green line is for samples with DOX, i.e. without signaling. For more details on AR calculation see methods. **h.** Elongation assay with B cells that activate p21 and CA-RhoA upon synNotch signaling activation. Left, schematic of B cell circuit. Right, micrographs of Day0 and Day6 of two conditions: top row, condition where synNotch signaling in B cells is allowed to happen; bottom row, condition where synNotch signaling in B cells is impaired with the small molecule Dox. In the micrographs the far-red signal from the constitutive far-red marker in A cells is rendered in blue, the mCherry signal in B’ cells is rendered in red, and the GFP signal is not shown. Scale bar is 500um. **i.** Graph of aspect ratio (AR) over time for n=2 +signaling experiments and n=1 no signaling experiment, each averaged from 3-6 technical replicates. AR =1 for a circle, AR>1 proportionally to the elongation of the ellipsoid. The continuous blue line is for samples with signaling; the dashed blue line is for samples with DOX, i.e. without signaling. For more details on AR calculation see methods. For more details on AR calculation see methods. Day 1 to 10 daily imaging time series and AR quantifications can be found in Fig. S8.

**Figure 7.**
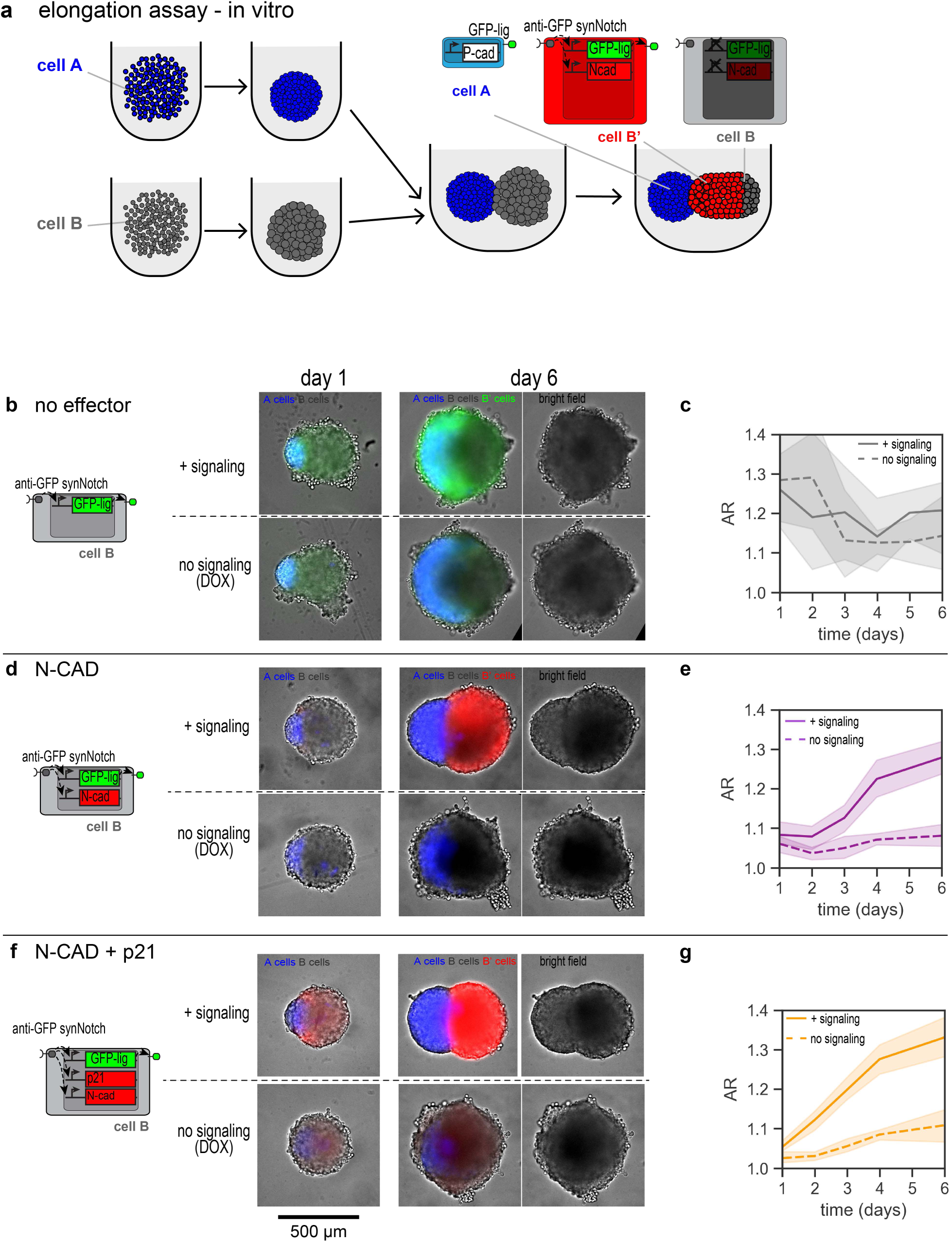
Programming tissue elongation through dynamical control of cell-cell adhesion and growth. **a.** Schematic of elongation assay: spheroids of 200 sender (A-type) cells (blue) and 4,000 transceiver (B-type) cells (gray) are seeded separately. B-type cells in this figure do not have constitutive N-cadherin. After 2 days, the two resulting spheroids are assembled in an individual well, and the assembloid is imaged daily henceforth. **b, d,** and **f.** For each row, on the left is a schematic of the gene circuits engineered in the transceiver cells. Sender and transceiver spheroids seeded as explained above were either cultured in doxycycline (dox, “no signaling” condition) or in control medium (“+ signaling” condition). Day 1 and 6 micrographs are shown for each condition, with miRFP703 represented in blue, GFP represented in green in b, and mCherry represented in red in d and f. The rightmost images are brightfield micrographs highlighting shape changes brought by activation of the synthetic gene circuit at day 6. **b.** Transceivers only induce GFPlig when activated. **d.** GFPlig, N-cad and mCherry are induced upon transceiver activation. **f.** GFPlig, N-cad, p21 and mCherry are induced upon transceiver activation. **c, e,** and **g.** Quantification of the aspect ratio (AR) over time of the conditions immediately on the left of each graph. See Methods and Figure S7 for more details on how the AR is measured. Briefly, the minimum AR that can be measured is 1 which corresponds to a perfect circle, any AR >1 corresponds to an elongated shape. Dashed lines represent AR measurements in the “no signaling” condition, and continuous lines measurements in the “+signaling” condition. The shaded regions represent the standard deviation from n=3-6 technical replicate in the n=1 experiment. Day 1 to 10 daily imaging time series and AR quantifications can be found in Fig. S10.

As characterized previously, the cells with the first class of circuits decrease both proliferation and fluidity in response to synNotch activation. Would a wave-like activation across a spheroid support tissue elongation, given the phenotypes observed *in silico* (Fig. 1b, S1c-d,f)? To answer this question, we performed elongation assays (**Fig. 6a**): a spheroid of 200 sender A-type cells is juxtaposed to a spheroid of 4,000 B-type cells, and their growth and signaling over time is followed with fluorescent microscopy. The choice of this ratio of cell numbers was made given that the measured difference of growth index in the A-type cells is roughly twice as high as that of B-type cells (with a DGR of 0.7 vs 0.35). To evaluate the impact of synNotch-dependent signaling, which supports the wave of activation, two media conditions were used: one where the signaling is allowed (no dox) and another one where signaling is completely abrogated (using the small molecule doxycycline that prevents TRE promoters’ activation). As shown in **Fig. 6b,d,f,h** in all cases, as expected, the “no signaling” condition resulted in rather spherical aggregates by day 6, and no or limited GFP or mCherry activation was visible in the B-type cell spheroids (see also **Fig. S8** for daily time points). We note that this is particularly significant here, as in these circuits the two spheroids express different cadherins P- and N-, which could induce partial establishment of greater-than-1 AR in other contexts, e.g. see **Fig. 4b-c** for similar experiments with different cell numbers). On the contrary, in conditions where signaling is allowed, mCherry is induced in activated B-type cells (thus generating B’-type cells). Whereas in the condition where B-type cells do not have effectors, this does not result in overt changes in the morphology, for B-type cells with effectors, the geometry of the aggregates is different in particular with a more visible demarcation of the A-type cells spheroid vs B/B’ spheroid. To quantify this effect with the different effectors, we measured the aspect ratio (AR=1 for a circle, >1 increasingly more elongated spheroid) of the system (see **Fig. S7** for details on the methods) over time, and plotted the results in **Fig. 6c,e,g,i**. These measures show that with signaling, the AR is increased over the 6 days already by p21 alone, and this increase is more pronounced with CA-MLCK and even more robustly with CA-RhoA. p21 induction alone resulted in the AR at 1.15-1.2 from day 1 to day 6 post-fusion, compared to a drop to 1.07 when signaling was inhibited in this same system. p21+CA-MLCK promoted an elongation of the system, from an AR of 1.15-1.2 on day 1 to 1.25-1.3 on day 6. p21+CA-RhoA promoted an even more robust elongation, reaching an AR of 1.3-1.35 at day 6. As shown in **Fig. S8e-j**, this effect is somewhat more limited in longer experimental times, except for p21+CA-RhoA. We also ran an elongation assay experiment with cell lines that induce effectors that were shown to have limited effects on fluidity and proliferation assays, i.e. constitutive N-cad and inducible p21, or inducible p21+KD-MLCK, inducible p21+Rac1DN. As shown in **Fig. S9**, as expected, fusion of sender spheroid with spheroids of these cell lines resulted in aggregates which maintained their aspect ratio across the 10 days of simulation without visible signs of increase.

Collectively, these results indicated that circuits capable of dynamically controlling tissue properties in space and time downstream of signaling achieve a dynamical structure whose AR is increasing over time for a certain period. The progressive spatiotemporal induction of genes controlling tissue growth and fluidity can promote tissue shape change, and genes that have a stronger impact on tissue fluidity and growth also promote stronger shape changes.

### *In vitro* implementation, complete circuits, ‘cadherin strategy’

Given that a flat and smooth B-B’ interface is important for elongation *in silico* (Fig. S1f), and induction of N-cad (via expression in only B’-type cells versus constitutive N-cad expression in both B-type and B’-type cells) resulted in a flatter signaling propagation front *in vitro* (Fig. 4j,k, Fig. S3f), we investigated a second class of circuits in which the B-type cells express N-cad downstream of synNotch signaling as an effector, either alone or in combination with p21 (**Fig. 7**). N-cad by itself was shown earlier to increase tissue growth and have little effect on tissue fluidity; whereas N-cad+p21 together, decreased tissue growth and had a trend for a marginal increase on tissue fluidity (**Fig. 5**). To test their effect on elongation assays, we fused senders A-type cell spheroid with B-type transceivers inducing N-cad or N-cad+p21 upon activation, at the same size ratios as in the previous assay (200 and 4000 cell spheroids, respectively) and monitored their AR after fusion (**Fig. 7a**). Here too, the dox-control showed limited or no mCherry (or GFP) activation, whereas the signaling conditions resulted in robust induction of mCherry (or GFP), indicating that the signaling system worked as expected to propagate synNotch activation from the A/B spheroids interface to the right-most end of the B/B’-type cell spheroids (**Fig. 7b,d,f**, see **Fig. S10** for daily time points). Although in controls where no effectors are induced signaling resulted in negligible changes in overall morphology (**Fig. 7b-c**), induction of the effectors was accompanied by a progressive increase of AR, which was reproducible over multiple experiments (**Fig. S10g**). While signaling-inhibited constructs have an AR measure around 1.0-1.1 across the 6 days of culture, transceivers that induce N-cad promoted an increase of the AR to 1.25-1.3 at day 6 and transceivers that induce NCAD + p21 promoted an even stronger increase of the AR to 1.3-1.35 at day 6. This peak aspect ratio stabilizes after day 6 on a plateau until day 10 (see **Fig. S10** for daily time points).

These results demonstrate that the induction of NCAD alone promotes the progressive increase of the aspect ratio of the system. Interestingly, the co-induction of p21 seemed to improve the elongation phenotype, potentially through inhibiting cell growth in the B’-type cells.

### Morphospace comparisons

Finally, we sought to contextualize *in vitro* and *in silico* results in a common “morphospace”, to help guide future efforts to improve the phenotypes we obtained through our *in vitro* work.

To do so, we first parameterized computational simulations to *in vitro* measures for parameters of proliferation and cell motility. Given that the description of the biological system in the *in silico* system is not specified at the molecular level, but at the higher agent-based level, we approached the parametrization in a “phenotypic” way: we performed parallel experiments *in silico* relative to the ones done *in vitro*, define and measure similar metrics, and then identified parameters *in silico* that resulted in phenotypic metrics close to the ones observed *in vitro*. Specifically, we first parametrized tissue growth *in silico*; *in silico*, cell proliferation is controlled by a parameter, “growth variable”, that controls how much a cell grows over time, since after a certain threshold of cell volume the cell is induced to divide. We established spheroids of *in silico* cells and had them grow for a time equivalent to 7 days, and calculated their daily growth rate, similarly to how we do it for the *in vitro* system (see Methods for details). We measured the growth index for a range of growth variable values and found the measured growth index to exponentially correlate with an increase in this growth variable (**Fig. S11a**); given that *in vitro* the growth index ranges between 0.5 and 0.1, we identified the range of the growth variable to generate that growth index to be between -4.5 and -6.

Secondly, we parametrized tissue fluidity. *In vitro*, this was measured via the 2 spheroids fusion assay (see **Fig. 5d-f**). We constructed a similar assay *in silico* too. The parameters that we thought could affect the rounding speed of a 2-spheroids system *in silico* are cell motility and cell-cell adhesion, so we performed a combined parameter search varying motility and cell-cell adhesion as indicated in **Fig. S11b**. *In vitro*, we measured a range of fluidity index between 10 (highly fluid) and 1um/h (more rigid). High values in this range were measured when both adhesion and motility were high (adhesion corresponding to homotypic N-cadherin as parametrized in^27^ for example, and motility at 40 units), and low fluidity was obtained when any or both of those were kept low: for any simulation with motility at 10, or cell-cell adhesion in the range of parental cells.

Finally, we asked, *in silico*, if proliferation changes would impact tissue fluidity, and if motility would impact tissue growth. We found that changes in proliferation do not affect the tissue fluidity readings for a range of cell motility (**Fig. S10c**); similarly, we noticed that a wide range of cell motility would not strongly influence tissue growth (**Fig. 10d**). In **Tables 1.4** and **1.5** are reported the computational parameters used for the parametrization scans. We notice here that some of these results diverged from *in vitro* results, especially the dependency of fluidity on cell-cell adhesion. This prompted us to re-evaluate fluidity modeling *in silico*. Upon further analysis, we noticed that the *in silico* system does not behave completely faithfully to the *in vitro* counterpart for the fluidity aspect (see more on that in the Discussion on the limits of the current computational model).

Altogether, with this parameterization, we identified *in silico* parameter ranges in this modeling framework that can be considered *in vitro* counterparts.

With this parametrized *in silico* system, we can now run simulations of elongation assays to explore the morphospace of phenotypes achievable with this range of parameters. To identify a space of maximal diversity of elongation assay outcomes, we decided to introduce the notion of morphospace (**Fig. 8**), inspired by recent work^77^. The first step towards creating a cogent morphospace for our system was to define meaningful morphological axes. The choice of the axes in any morphospace is both very consequential for the interpretation, and, to a certain extent, arbitrary. For this specific choice, we were guided by the computational sensitivity analysis done for **Fig. 1**, and by the study of the literature of tissue fluidity and elongation. The choice for the 3 axes was as follows: in x, an index of the ratio of growth index between B’-type and B-type cells; in y, a measure of the resistance to rounding (that is, the opposite of the fluidity index) of tissues made of B’-type cells; and in z, a measure of the isolation of the B-type cells from the B’-type cells. These parameters were shown from the sensitivity analysis to be important for contributing to elongation separately. To generate numeric values for these quantities, the x-axis is the ratio of proliferation index in B vs B’-type cells; the y-axis is measured as 1/tissue fluidity index of B’-type cells; and the z-axis is a qualitative measure of isolation of B-type cells from the B’-type cells.

**Figure 8.**
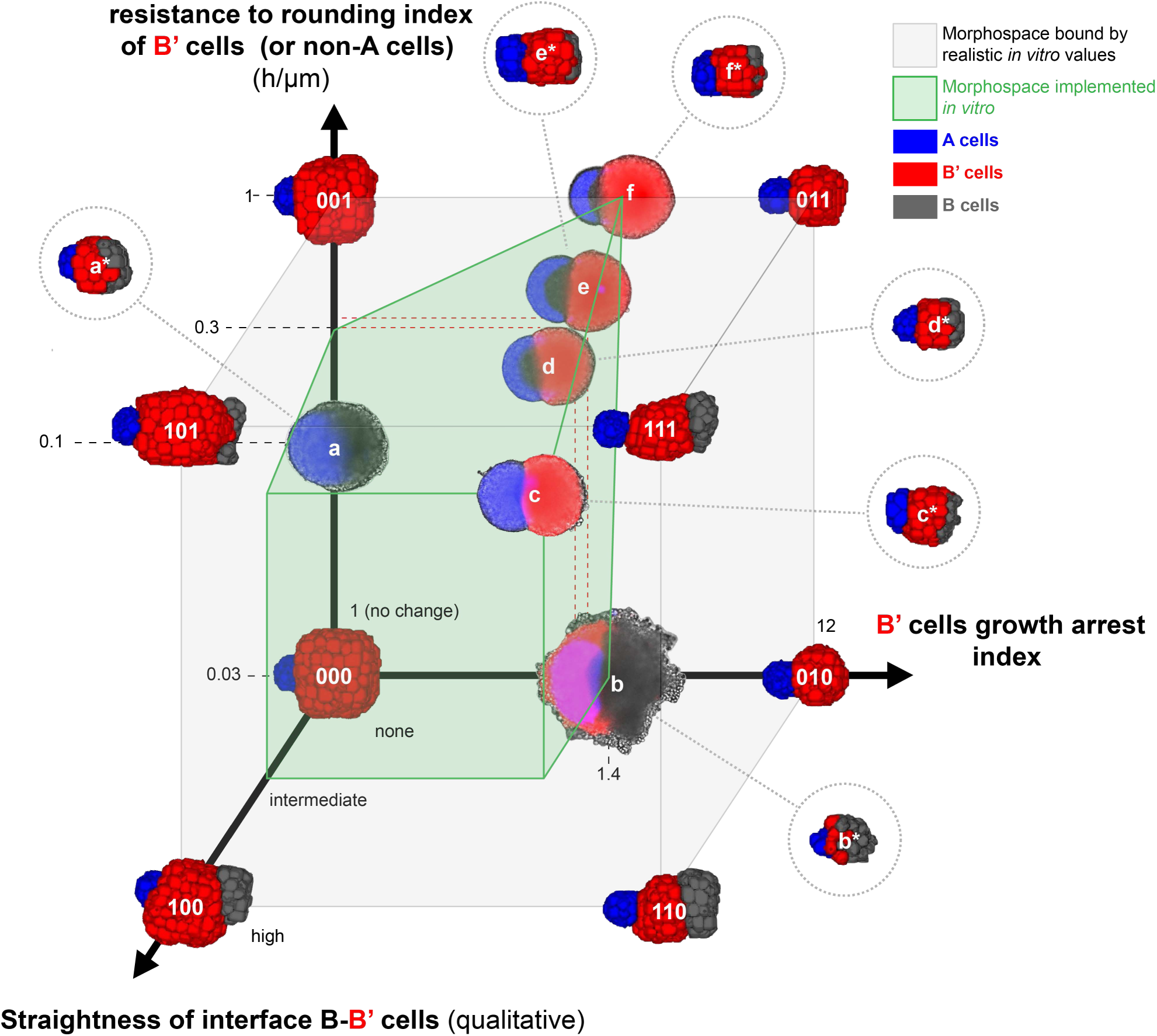
A common morphospace for *in silico* and *in vitro* results. **a.** 3D Morphospace graph. Throughout the graph, A-type cells are in blue, B-type cells are in gray, and B’-type cells are in red. The axes are as follows: x-axis, the ratio of growth index in B vs B’-type cells (which is proportional to B’-type cells growth arrest); y-axis, 1/tissue fluidity index of B’-type cells, which is a measure proportional to B’-type cell spheroid viscosity; and z-axis, a qualitative measure of the isolation of the B-type cells cluster from the B’-type cells cluster (correlating with low heterotypic/high homotypic adhesion of B and B’-type cells). The light gray box is bounded by the highest value which can realistically be achieved from each axis according to our *in vitro* data. At the 8 vertices of this white cube are placed the corresponding endpoint (7 days) elongation assay snapshots from *in silico* implementations. Corresponding circuits were numbered as in this figure, from 000 to 111, in Table 1.2. The green bounding lines define the volume in the morphospace that contains implementations that have been realized *in vitro*. On the bounding lines are shown selected examples of *in vitro* realizations, from day 6 micrographs where the resulting spheroid structures are isolated from the background via image processing. They go from a-f. For all of them, A-type cells have GFP ligand and P-cadherin expression, and spheroids were seeded from 200 cells before fusion. B-type cells were engineered from the “fast” clone chassis, and expressed the following constitutive of inducible effectors: a: constitutive N-cad, no inducible effectors (and thus no induced mCherry), also shown in Fig. 6b; b: inducible p21, also shown in Fig.S10a; c: inducible N-cad+p21, also shown in Fig. 7f; d: constitutive N-cad and inducible p21, also shown in Fig. 6d; e: constitutive N-cad, inducible p21+CA-MLCK, also shown in Fig. 6f; f: constitutive N-cad, inducible p21+CA-RhoA, also shown in Fig. 6h. For each one of these are also shown the corresponding *in silico* implementation (a*-f*, circled by a dashed line), which in the graph are connected to the corresponding *in vitro* implementation with a dashed line. For the complete *in silico* genomes for a*-f* See Table 1.3. See text for further description. *In silico* – day 7 for the vertices; for the asterisks is variable, see Fig. S12a, last column.

We first placed *in silico* results in this morpho space for simulations that have maximal values of the parameters, within the identified bounds of reasonable biological implementation (**Table 1.2** for parameters used in these simulations). The *in silico* genotype that minimizes all three axes (000) is obtained with parameter values that have B-type cells behave like parental L929 cells constitutively expressing N-cad: B-type cells still convert to B’-type cells, but both B-type and B’-type cells share the exact same features, so nothing changes in the B/B’ side over time. We then defined ways to move along the 3 axes defined above. To increase B’-type cell growth arrest, i.e. the x axis, we tuned the “growth variable” parameter. To increase the resistance to rounding of B’-type cells (y axis) to the maximum, we decided to tune the motility of B’-type cells. To increase the z axis, we decided to decrease B-B’ adhesion, which was shown in the sensitivity analysis to affect interface shape. (**Fig. 1**)Then to traverse the space outside of each axis, we linearly combined the parameters. We showed that there was no significant crosstalk between fluidity and growth *in silico*, and assumed that changing the growth index and/or the fluidity index of B’-type cells would not significantly affect the B’-B interface shape. As shown in **Fig. 8**, maximum elongation was achieved when the 3 axes were maximized: high separation between B-B’-type cells, strong reduction of growth in B’-type cells compared to B-type cells, and high resistance to rounding of B’-type cells.

We then placed *in vitro* implementations (and their computational counterparts) in this same morphospace (See **Table 1.3** and **Fig. S12a** for parameters used for these simulations). We found that the *in silico* realizations of *in vitro* implementations shared some aspects with the *in vitro* results but diverged in others. In both *in vitro* and *in silico* elongation experiments, A-type cells were confined on a pole on the left of the structure, and B/B’-type cells were on the opposite pole and, when present, inactivated gray B-type cells were on the opposing pole compared to the blue A-type cells. Moreover, the global aspect of the collective structure is more spherical for structures closer to the origin of the axis and becomes more “elongated” moving away from the origin and towards the 1.1.1 point that maximizes the 3 axes. On the other hand, we observe that the *in silico* implementations tend to be more compacted and slimmer compared to the *in vitro* ones (e.g. e/e* or f/f* in **Fig. 8**). This results in achieving a higher aspect ratio *in silico* compared to *in vitro* and might be linked to the limitations of the *in silico* model (see Discussion).

We also noticed that the *in vitro* implementations covered a smaller portion of the entire morphospace (green box in the morphospace in **Fig. 8**). Given that the amount of elongation in general in the morphospace is proportional to their proximity to the max-elongation corner, the design indication for maximizing elongation would be to maximize these 3 axes. This provided the basis for our last design: the fact that the z-axis is crucial for elongation in the parameterized version of the morphospace led us to design a still untested circuit that would maximally support elongation as follows: B-type cells would express N-cad and, in response to synNotch activation, they would switch to P-cad, i.e. would repress N-cad and start expressing P-cad (**Fig. S12b,c)**. This would maximize this axis in a future implementation *in vitro*.

Collectively, these analyses suggest that *in vitro* elongation with the current circuits is sub-optimal compared to the *in silico* counterparts and could be improved by generating cells with stronger control of rigidity and better B-type cell confinement effectors.

## Discussion

Identification of novel circuits for synthetic morphogenesis and their implementation in mammalian cellular systems are still bottlenecks for the application of a synthetic morphogenesis approach based on programmed self-organization. Here, to widen this bottleneck, we show our approach to programming the axial elongation of a spheroid of mouse embryonic fibroblasts with synNotch-based genetic circuits. This pipeline encompasses *in silico* circuit identification, *in vitro* development of circuit parts development and assembly in complete circuits guiding tissue elongation. What have we learned?

Designing and testing circuits through a computational model gave us great freedom and lowered the entry barrier for exploring and testing multiple approaches quickly. The model allowed us to perform sensitivity analyses which guided our search for effector genes, even though it was partially parametrized. The system parametrization allowed us to generate a morphospace where we contextualized both *in vitro* and *in silico* implementations (see more below), thus creating an *in silico* analogue of the *in vitro* implementation, following a paradigm of computational design environment common in engineering disciplines. While issues exist in the specific model we used (see below), we believe that the path charted here can be inspiring for future attempts to identify circuits for tissue self-organization: computational tools could be used to not only explore critical design parameters, but also to design the circuits themselves. The space of possible circuits is much larger than the parameter space of a single circuit; exhaustive searches as the ones deployed in the past for intracellular polarization circuits^78^ may not be possible; machine learning-based approaches or other algorithmic methods^79^ will be required to intelligently explore this space.

When choosing a computational platform we were confronted with a difficult balance between 1) models that properly simulate tissue physics but are so computationally intensive they are often used in a 2D setting, such as Voronoi or vertex-based models, and 2) simplified models that have less parameters and are faster to run in 3D. We settled on using a model of the latter category: the CC3D-based GJSM model. This model has been previously reported^27^ and is modified to model gene circuits and parametrized synNotch signaling. We made this choice to maximize parametrization and runtime efficiency, in order to quickly screen multiple configurations to inform *in vitro* work.

For parametrization, given the limited number of parameters in this system, we decided to use a “same experiment” approach, where analog assays were performed *in vitro* with biological cells and *in silico* with computational equivalent cells. Tuning one parameter (e.g. the proliferation-controlling growth parameter) to match the observation (e.g. the measure of the growth index) *in silico* vs *in vitro* allowed us to proceed with qualitatively accurate parametrizations.

For managing computational simulations duration, we reduced the number of cells *in silico* by one order of magnitude, from thousands of cells *in vitro* to 100s of cells *in silico*. Whether we can more faithfully simulate in vitro phenotypes by using a larger number of cells in silico is an important question, which will require increased computational power for its exploration.

Additionally, the dependency of the system on the initial number of each cells is an interesting parameter which could be further optimized by in silico and in vitro screens.

While the model we used revealed useful for running multiple simulations in parallel, we noticed it was limited in its mechanic realism. Indeed, although CC3D is commonly used for implementing cell-cell signaling, adhesion and proliferation phenomena, in addition to simulate some already elongated structures^80,81^, we noticed a discrepancy from in vitro results about tissue rounding/viscosity. Simulated non-spherical tissues past a certain size *in silico* would not as readily round up as the *in vitro* counterparts. We attempted to modify parameters in the simulation, such as cell motility, adhesion, the parameters of the “medium” cells (the black space surrounding the “tissue”), and the initial conditions, but these did not solve this issue. Such a problem could be in our simulation case partly circumvented with the available tools in CC3D by adding a centripetal attraction field. More broadly, the field is currently lacking a computational 3D tissue model in which tailored signaling can be implemented in parallel with accurate tissue physics. Future algorithms will probably be based on machine learning approaches, as evidenced by the recent paradigm-breaking efficiency of AlphaFold in predicting protein folding.

From this initial computational approach, we implemented our design *in vitro*. There, we found it helpful to first breakdown the circuit into modular subroutines such as signal propagation, signaling-mediated growth control, and signaling-mediated tissue rigidity control. This proved to be a good method for designing synthetic morphogenetic circuits, dividing the overall design task and implementation into more manageable optimization problems. With this approach we were able to identify effectors relevant to efforts to control tissue physics in the future, as both proliferation control and tissue fluidity control were achieved here for the first time downstream of a synthetic signaling pathway. Interestingly, we obtained viscocapillary speed (“fluidity index”) values in the range of 1 to 10 µm/h. While the range is comparable to results reported with spheroids made from various cells^50,82^, the combined induction of p21 and CA-RhoA expression resulted in an average VCS of 1.2 µm/h, and an all-time minimum of 0.9, which is to our knowledge one of the lowest values reported through this spheroid fusion assay. This suggests that the combination of those two effectors could be useful in future efforts to increase the viscosity of mouse embryonic fibroblast tissues. We want to note that here we were testing individual effectors, but the flexibility given by synthetic circuit design would allow us to combine multiple effectors, and even non-natural ones, to further improve the control of cell behaviors.

We then moved on to generate complete circuits for elongation assays. While we broke up our engineering efforts into specific modules, we did not follow a completely exhaustive implementation of complete circuits but favored a kind of “additive design” downstream of two main branches of implementation. The two branches are defined as based on whether they use constitutive or inducible N-cadherin in B-type cells. Within these branches, we would screen effectors in a context that would be relevant to the previously designed part, so that the interaction of those modules would already be accounted for in the results of the assay. For example, the impact of the expression of proliferation effectors was assessed in the context of the transceiver circuit. Also, the impact of the fluidity effectors was studied conjointly with the induction of the proliferation effector we retained from the previous step (p21). On one hand, this allowed for a faster transition from effector screen to test in elongation assays which allowed to test all the effectors in elongation assays, even the ones that were not performing highly in the growth or fluidity assays. These generated interesting negative results for the elongation assays themselves.

The balance between how much one can do a combinatorial approach vs a stepwise building on previous successes lies on the throughput of the cell line engineering and on the throughput of the assays. Increasing these would allow us to perform more exhaustive searches and combination of circuit space. This increased screening power would permit to identify individual effectors for efficient control of the parameters of interest, and would also open the way to two promising screening approaches: 1) the screening of larger number of individual effectors, either natural, synthetic or mutated from natural; 2) the systematic combination of effectors targeting a single phenotype, which could reduce off-target effects while increasing on-target efficiency.

The two main design strategies we exploited to guide tissue elongation demonstrated different degrees of control of aspect ratio over time. In the first strategy, B-type cells constitutively express N-cadherin and induce either p21 or fluidity effectors CA-MLCK or CA-RhoA. This supports the notion that changes in cytoskeletal tension is an excellent tool for guiding tissue morphogenesis. Specifically to this example, the induction of a wave of of constitutive active forms of MLCK and RhoA generated a strong separation between spheroid of sender A-type cells, and the activated B’-type cells, and seem to increase resistance to rounding, as previously proposed^83^. CA-RhoA revealed particularly efficient in this regards, which was our strongest effector identified in the screenings for viscosity modulators. Accordingly, effectors that induced limited changes in fluidity index generated spheroids that did not change their AR over time, thus validating the importance of lowered rounding speed for guiding tissue elongation. It is to be noted that the AR increase that is obtained with this strategy in the first 5-6 days subsides in the following days. This could be due to a limitation of this strategy past a certain system size which is reached only after day 6; or one that could be achieved with increased proliferation control.

In the second strategy, the B-type cells do not express cadherin in the basal state but activate in a wave fashion the effector N-cadherin, either alone or alongside p21. In elongation assays performed with these cells, the increase of AR over time is even more dramatic and does not revert to basal even after 10 days. Although this is somewhat expected from the fact that constitutive heterotypic adhesion supports the formation of dumbbell shape structures within a certain cell number range,there seems to be something more at play here. Indeed, all the B-type cells of the first strategy (above) constitutively express N-cad, but N-cad expression only does not lead to AR increase over time, and even the addition of strong fluidity control effectors result in decreasing AR over long time course. This suggests that including the expression of a cadherin in a wave-like fashion can be a robust way to generate stable domains in growing spheroids or assembloids. This strategy could be interesting to port in the context of stem-cell-derived organoids and assembloids, to increase the separation between single organoids^84^. It remains to be explored whether these results are dependent on initial number of cells, as some of our previous computational results seem to suggest for other phenotypes.

Finally, we find it very helpful and instructive to contextualize *in vitro* and *in silico* achievements within a common morphospace. It helped us rationalize relevant axes, locate implementation and compare *in silico* with *in vitro* results; additionally, it provided a sense of how distant the *in vitro* and *in silico* implementations were, and provided a basis for the design of the next generation of circuits. We also imagine a version of the morphospace with more dimensions, where embedding approaches such as UMAP or t-SNE could reveal interesting for visualizing many results on an n-dimensional space (each dimension a parameter of the circuit) according to their graph-level proximity, as was done elsewhere for visualization of gut organoids phenotypes^85^.

The morphospace visualization also suggested that stronger elongation phenotypes are not so distant from the best results we achieved *in vitro* along those axes. Why were we not able to reach results similar to the ‘upper corner’ implementation *in silico* (111) that maximizes the three axes and can anything be done in the future to improve on this initial prototype? This question has different answers for each axis as follows.

For the x-axis (growth control), the *in silico* genome allows us to use the most extreme values of basal B vs induced B’-type cell proliferation, with a growth index drop from 0.6 to 0.05. Although both values of DGR are observed *in vitro*, we were not able to generate a cell line that is able to maintain a basal of 0.6 and an induced value of 0.05. Even for a B-type cell that induced p53, our strongest proliferation inhibitor, the basal, uninduced proliferation is lowered to around 0.4, probably due to leakiness of expression of effectors; this explains why we cannot have maximal *in vitro* implementation along this axis. To maximize this axis, a better gating of the on/off induction via synNotch would be needed. Simultaneously inducing multiple effectors targeting tissue growth could lead to better and more specific phenotype control.

For the y-axis (fluidity index), similarly, the *in silico* genome allows us to use the most extreme values of basal B vs induced B’ viscocapillary speed measures of 20 to 0.11. Again, these are observed *in vitro*, but we still cannot switch from 20 to 0.11 in a synNotch-gated manner. The strongest effectors in fact go from 4.5 to 2.5 or from 2.6 to 1.2 on average. A larger delta, with un-lowered basal, would allow us to maximize this axis *in vitro*.

Additionally, for these two axes combined, we noticed that the best effectors for decrease of proliferation (p53) affected viscosity in the unwanted direction (increased fluidity), which prevented us to maximize both axes concurrently with this effector. Search for improved proliferation effectors or combination thereof should keep the optimization of both parameters in mind.

Finally, for the z-axis (B-B’ interface), the maximal possible straightness of the interface between B and B’ is obtained when their heterotypic adhesion is kept low. In our *in vitro* implementations this axis is increased by having cells express or not cadherin molecules, which leads to intermediate heterotypic adhesion between B and B’. To obtain lower B-B’ heterotypic adhesion values, these cells should express cadherins with low heterotypic strength, like the pair P and N, which has been identified in our improved design which obtains this through a NOT gate downstream of synNotch to decrease P-cad expression upon activation, while N-cad expression would be directly increased upon receptor stimulation.

One final point for discussion is how much the synthetic circuit-based approach to synthetic morphogenesis stakes compared to other potential methods of controlling self-organization. In the larger field, whether the genetic circuits are the “best” layer for intervention is debated^14^. We think that it is good for testing biological hypotheses on genes and cell behaviors underlying complex biological phenomena; it could emerge as an independent, more basic branch of developmental biology. For building purposes, it gives the impression that we are acting at a very basic layer, with genetic circuits, while the emergent properties are multiple layers away, and as such hard to control directly. So maybe a combination approach could get the benefit of all the worlds?

Globally, our efforts highlight that using synthetic gene circuits built from the ground up is a very challenging approach past a certain complexity threshold, as the tools for standard engineering are not yet completely adapted to the engineering of noisy, soft, extremely complex biological systems with their own pre-evolved self-organization modules. We attempted to work with pre-existing modules at the level of effector genes; an alternative would be to invoke higher-level morphological modules towards the target goal plasticity and train it toward a target goal, as was proposed elsewhere^14^. A further alternative would be a combination approach of both entry-points: high-level rules for control of morphological plasticity, coupled with lower-level control via genetic circuit for finer and more freely user-designed developmental and morphogenetic programs.

Collectively, this work represents a further step towards the rational design of synthetic genetic circuits for morphogenesis, a step closer to the dream of rationally designing and implementing developmental mechanisms to program new self-organized morphologies and tissue architectures.

## Methods

### Constructs

The synNotch-encoding plasmid used to generate the transceiver cells is available on Addgene (pHR_SFFV_LaG17_synNotch_TetRVP64, catalog number #79128). The GFPlig-encoding plasmid used to generate the transceiver cells is available on Addgene (pHR_EGFPligand, catalog number #79129).

Mouse *WNT5A* was amplified from plasmid pRK5-mWnt5a [bought from Addgene, catalog number #42279]; human CA-RhoA (a constitutively active mutant of *RHOA, RHOA^Q63L^*) was amplified from plasmid pRK5-myc-RhoA-Q63L [bought from Addgene, catalog number #12964] *CA-MLCK* (constitutively active mutant of human *MLCK, MLCK^ED785-786KK^*) was amplified from plasmid pSLIK CA MLCK [bought from Addgene, catalog number #84647]; *KD-MLCK* (kinase-dead mutant of human *MLCK, MLCK^E1626K^*) was amplified from plasmid Hela kinase-dead MLCK-GFP [bought from Addgene, catalog number #46317]; human *CDKN1A* (p21) was amplified from plasmid Flag p21 WT [bought from Addgene, catalog number #16240]; human *CAV1* (Caveolin-1) was amplified from plasmid mCherry-Caveolin-C10 [bought from Addgene, catalog number #55008]; mouse *SOX9* was amplified from plasmid tetO.Sox9.Puro [bought from Addgene, catalog number #117269]; human *RAB5A^DN^* (dominant negative form of *RAB5A, RAB5A^S34N^*) was amplified from plasmid mCherry-Rab5DN(S34N) [bought from Addgene, catalog number #35139]; human *CDKN2A* (P16INK4a, p16) was amplified from plasmid pQCXIH-CDKN2A [bought from Addgene, catalog number #37104]; human *CDKN1B* (p27) was amplified from plasmid pcDNA3-myc3-p27 [bought from Addgene, catalog number #19937]; human *TP53* (tumor protein 53, p53) was amplified from plasmid pLenti6 / V5-p53_wt p53 [bought from Addgene, catalog number #22945]; *mCherry* was amplified from plasmid pHR TRE3GS->mCherry_pGK->BFP [bought from Addgene, catalog number #162231]; human *LOXL2* was amplified from plasmid pQXCINeo-LOXL2-BC [bought from Addgene, catalog number #134763]; human *TG2* was amplified from plasmid W118-1_hTGM2-Flag [bought from Addgene, catalog number #180407]; human *Cdh1* (ECAD) was amplified from plasmid E-cadherin-GFP [bought from Addgene, catalog number #28009]; mouse *Cdh2* (NCAD) was amplified from a plasmid given by the Lim lab^21^; human *Cdh3* (PCAD) was amplified from plasmid pcDNA3 P-cad [bought from Addgene, catalog number #47502]; H2B-miRFP703 was amplified from plasmid pH2B-miRFP703 [bought from Addgene, catalog number #80001].

Plasmids were generated using the In-Fusion Cloning Kit (Takara Bio). For constitutive expression, fragments were inserted in a pHR backbone with an SFFV promoter. For synNotch-mediated induction of expression, they were inserted in a pHR backbone downstream of a TRE promoter, aligned with the following downstream coding sequence: P2A-mCherry, and followed either by PGK>tagBFP for FACS sorting of positive cells or PGK>PuroR / PGK>HygR for antibiotics mediated selection.

### Cell culture

L929 cells (ATCC #CCL-1) were cultured in DMEM (High glucose, +L-glutamine, -Sodium pyruvate, Thermo Fisher Scientific) supplemented with 10% FBS (Qualified, US origin, Thermo Fisher Scientific) and 100 U/mL Penicillin / Streptomycin (Thermo Fisher Scientific). Passaging was performed by a DPBS (Thermo Fisher Scientific) wash followed by TrypLE Select (Thermo Fisher Scientific) cell dissociation. Transceiver cell lines were routinely passaged with 0.1 mg/L tetracycline to keep them in the OFF state.

### Cell line engineering

All cell lines were engineered by lentiviral transduction followed by antibiotic selection or Fluorescence Assisted Cell Sorting (FACS).

Lentiviral vectors were produced by transfecting Lenti-X™ 293T cells (Takara Bio) with the plasmids psPAX2 [Addgene catalog number #122]6) and pMD2.G [Addgene catalog number #12259] in combination with the plasmid containing the cassette of interest inserted in a pHR backbone, using the Lipofectamine LTX transfection kit (Thermo Fisher Scientific). Culture medium containing lentiviral particles was harvested 2 to 3 days following transfection, sterile filtered using a 0.22 µm pore size polyethersulfone (PES) filter (Genesee Scientific), and finally purified and concentrated in DPBS using the Lenti-X Concentrator kit from Takara Bio. Cell lines lentiviral transduction was performed by exposing the cells to a range of concentrations of lentiviral particles for 3 days, before triple washing with DPBS and passaging to a new culture plate.

FACS was performed on the healthy surviving cells that had been exposed to the highest lentivirus dose using a BD FACSAria IIIu cell sorter.

Transceiver cells were generated from parental L929 using constructs pHR_SFFV_LaG17_synNotch_TetRVP64 and pHR_TRE_GFPlig. Cells were sorted based on myc staining (myc tag located in Nter of the synNotch CDS) (anti-myc antibody 2233S from Cell Signaling Technologies) and GFP signal. Individual cells were sorted to generate monoclonal populations. Clone T2 and T3 were selected based on spheroid formation ability and 3D signaling speed when fused to a sender cell spheroid.

All other generated lines were polyclonal.

Sender cells were generated from parental L929 using constructs pHR_SFFV_GFPlig and pHR_SFFV_H2B-miRFP703. Cells were sorted based on GFP and miRFP703 signal. Pcad expressing senders were obtained using construct pHR_SFFV_PCAD, followed by sorting of cells stained by an anti-PCAD antibody (Santa Cruz #sc-74545).

Transceiver cells with inducible effectors were generated by transducing “fast” transceiver cells with lentiviral vectors, resulting in the insertion of the cassette containing TRE3G>*gene_of_interest*-P2A-mCherry followed by either PGK>PuroR, PGK>HygR or PGK>tagBFP. Polyclonal populations were selected either by puromycin selection with 20 mg/L for 3 days, by hygromycin selection with 400 mg/L for 6 days, and/or by sorting of tagBFP-expressing cells.

### GFP production + purification

GFP was purified as an N-terminal hexahistidine fusion protein. To express GFP, BL21(T1R) E. coli cells were grown to an optical density of 0.5 from an overnight-grown glycerol stock, chilled to 25 °C, induced with 1 mM IPTG (Sigma, I6758), and allowed to express for 5 h. GFP was purified by NEBExpress Ni Spin Columns (New England Biolabs, S1427S) following the manufacturer’s instructions, dialyzed against 1× PBS overnight at 4 °C, sterile filtered, and frozen at −80 °C until use.

### Spheroids generation

Cells of interest were counted using a Countess II (Thermo Fisher). Cell suspension was seeded in a non-adhesive U-bottom 96-well plate (Nunclon Sphera, Thermo Fisher Scientific)). Culture plate was centrifuged for 2 min at 100 g. Cell number was then evaluated by eye and corrected if necessary. Cells were left for 2 days to coalesce in a spheroid before handling.

### Spheroids fusion

Spheroids of interest were fused using a multichannel micropipette. Briefly, some medium was pumped in the tip, then quickly ejected to displace the spheroids from the bottom of the well. All the content of the well was then pumped in and transferred to a well of interest containing the spheroid to be fused with. Culture plate was centrifuged for 1 min at 50 g to bring the 2 spheroids in close contact before imaging. For signal propagation experiments with senders and transceivers, the transceiver spheroids were seeded in tetracycline 0.1 mg/L. After 2 days, the transceiver spheroids were gently rinsed several times to remove tetracycline before being brought in contact with the sender spheroids.

### Imaging

Imaging was performed using a BZ-X710 (Keyence) fitted with a CFI Plan Apo 10X objective and a Tokai Hit live imaging setup.

### Image analysis

Image analysis was performed using Fiji^86^. Data analysis was performed using Python, Jupyter Notebook, and GraphPad Prism.

### Measure of signal propagation rate in transceivers

1000-cell spheroids of transceivers of interest and of senders were seeded as described above. After 2 days, transceiver spheroids were rinsed as described above, and spheroids were brought together and imaged every hour. At every 12h time increment, the distance between the sender-transceiver fusion border and the activated transceiver propagation front was measured at 2 GFP signal thresholds, either on the symmetry axis going through the center of both spheroids, or at the maximally distant point from the spheroids fusion border. Indeed, the propagation front in the center of the transceiver spheroid lagged propagation on the borders. Those 4 measurements were then averaged to obtain the distance used for plotting. This quantification method allowed us to consider the central and lateral components of signal propagation speed.

### Fate switch in transceiver cells by culture on GFP-coated plate

An *in vitro* assay was devised to evaluate the impact of various genes of interest (GOIs) on tissue growth and fluidity. “Fast” transceivers were engineered with pHR plasmids containing the TRE>*GOI*-P2A-mCherry cassette. Resulting transceivers with inducible effectors were either activated (ON, GOI overexpression) by culture for 3 days on a GFP-coated plate (see below), without tetracycline or doxycycline, or inhibited (OFF, minimal GOI expression) by culture for 3 days on a standard plate with 10 mg/L doxycycline. Because of lateral signaling, spheroids made from ON transceivers stayed activated for the duration of the experiment, while spheroids made from OFF transceivers remained inhibited by the continued presence of 10 mg/L doxycycline.

GFP-coated plates were prepared as follows: 48-well plates (Genesee Scientific) were filled with 100 µL of purified GFP diluted at 0.1 mg/mL in distilled H2O and left to dry open overnight under a biosafety hood.

### Measure of volumetric growth rate (growth index)

1000-cell spheroids of ON or OFF transceivers of interest were seeded as described above. Spheroids were then imaged at a minimum on days 2 and 7 post-seeding. Ellipsoids were manually fitted to best approximate the shape of the spheroids. Spheroid volume was computed by using the oblate ellipsoid volume formula. Daily growth rate (growth index, DGR) was then computed using the following formula:

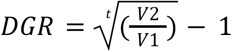

with *t* the elapsed time in days between t1 (2 days post seeding) and t2 (7 days post seeding), *V1* and *V2* the volume of the spheroid at t1 and t2, respectively. A growth index of 0 means no growth at all; a growth index of 1 means that the volume doubles every 24 hours.

### Measure of viscocapillary speed (fluidity index)

Viscocapillary speed was measured in the same fashion as in^50^.

Briefly, 1000-cell spheroids of ON or OFF transceivers of interest were seeded as described above. 2 days after seeding, spheroids were fused and centrifuged at 50g for 1 minute. Spheroid pairs were then immediately imaged every 30 min in Brightfield for 24 hours. Using Matlab, spheroids were binarized with an Entropy threshold. Geometrical descriptors (aspect ratio, theta angle between spheroids contact border and spheroids centers) were extracted. Whole-system relaxation rate was measured from the decay of the theta angle or the aspect ratio over time. Viscocapillary speed (fluidity index) was then computed from the whole-system relaxation rate by correcting for spheroids volumetric growth rate.

### Measure of mCherry fluorescence level

mCherry fluorescence levels were measured in 2-day-old spheroids formed from transceivers cultured either on a GFP-coated plate or on a standard plate in the presence of 10 mg/L doxycycline. Measurements were made from epifluorescence micrographs. A region of interest encompassing the whole spheroid was drawn from the brightfield channel of the micrographs, either automatically shaped with the Find Edges algorithm of Fiji or manually traced. The average signal intensity was then measured in the region of interest in the mCherry channel. Normalization was performed in each experiment by dividing all results with the average intensity obtained in spheroids made of OFF transceivers with no effectors.

### Flow cytometry

Analysis of mCherry and GFPlig levels at the single cell level was performed by Flow Assisted Cell Sorting, using a FACSAria II (Beckton-Dickinson) followed by gating with the FlowJo software and analysis with Python. Cells were dissociated, washed and processed after 3 days of culture either on a GFP-coated plate or on a standard plate in the presence of 10 mg/L doxycycline.

### Measure of aspect ratio (AR)

Analysis was performed from brightfield micrographs. Contours of cellular aggregates were obtained either manually for difficult cases, or with a Fiji automated image analysis pipeline for most: 1) Find Edges; 2) 20x Smooth (Gaussian Blur); 3) Thresholding; 4) Fill Holes; 5) Keep Largest Region (MorphoLibJ plugin). We then used Fiji Measure Particles to obtain the minor and major axes of the best-fitting ellipse. The ratio of those yields the aspect ratio (AR) of the shape, with an AR of 1 corresponding to a perfect sphere.

### Computational model

We utilized CompuCell3D (CC3D) 4.1.1 to computationally model cell behavior through simulations of cells using the cellular Potts formalism as described in the GJSM model^27^. CC3D contains features that replicate multiple aspects of *in vitro* cell behavior, which were modified using CC3D Python v3.7.7 according to manual v4.1.2.6. To maintain simulations consistent with *in vitro* assays, the Gaussian distribution for the assignment of the initial cell target radius was modified (µ = 3 pixels, σ = .425 pixels) from the distribution used in^27^. 100×100×100 and 120×120×120 simulation lattices were used. Cubes of sender cells, transceiver cells, or different cubes of both were placed in the center initially. Cells rapidly formed sphere-like structures, consistent with *in vitro* initial conditions.

### *In silico* control and measure of viscocapillary speed (fluidity index)

For parameterization, the viscocapillary speed (fluidity index) was measured by using two initialized spheroids of sender cells with approximately 64 cells each at the beginning of the simulation. Viscocapillary speed was modified by altering the basal motility variable in the code. The inputs of this variable range from 0 to 100, where 0 represents no motility and 100 represents extremely motile behavior. Data was collected for 48000 MCS to represent 2 days’ worth of real-time data. In elongation simulations, initial sender and transceiver cells were set to have approximately 64 cells at the beginning of the simulation. Each cell type was given its own respective basal motility value. The spheroid size was measured and quantified as described in the *Measure of viscocapillary speed* section (Grosser et al., 2021).

### *In silico* control and measure of volumetric growth rate (growth index)

For parameterization, individual growth rates (growth index) were measured with an initialized spheroid of sender cells with approximately 64 cells at the beginning of the simulation. The proliferation/volumetric growth rate of spheroids was controlled using a MitosisSteppable class where the sender cells would grow into a specific target radius depending on the MTFORCEMAX and MTFORCEMIN values provided. MTFORCEMAX indicates the positive mitosis drive force fluctuation, set at a value of -3 x 10^(-1*X), X being the adjusted numerical value. MTFORCEMIN indicates the negative mitosis driving force fluctuation, set at a value of 4 x 10^(-1*X). X values are manually edited in the code and the same value is set for both the MTFORCEMAX and MTFORCEMIN variables. As the exponent values for both variables become more negative, the growth rate of the spheroid slows; a more positive exponent value increases this rate. When the individual cells reach a threshold volume the cells divide into two. Data was collected for 168000 MCS to represent 7 days’ worth of real-time data. For elongation simulations, the initial sender and transceiver cells were set to have approximately 64 cells at the beginning of the simulation. Each cell type was given its own respective MTFORCEMAX and MTFORCEMIN values; cells were removed entirely from the MitosisSteppable class if they were intended to not have proliferation during the specific experimental simulation.

Spheroid size measure and calculation of growth index were performed similarly to *in vitro* (fitting of an ellipsoid on 2D snapshots of the spheroid, calculation of volume with oblate ellipsoid formula, calculation of daily growth rate).

## Authors contribution

CL realized the first *in silico* implementation of the circuit for axial elongation. JC led the experimental section. CC, NP and CS generated the *in silico* results. JC, CC, CS and NJ generated the constructs, engineered cell lines and generated *in vitro* experimental data. JC designed the experimental and data analysis protocols. JC, CC, CS and NJ performed data analysis. SG computed the viscocapillary speed from homotypic spheroids fusion data. LM acquired the funding for the project. LM and JC managed the project and wrote the first draft of the manuscript.

## Acknowledgements

We thank the Flow Cytometry Facility and Optical Imaging Facility of the Eli and Edith Broad CIRM Center; the Morsut lab for their scientific input and support; Matt Thomson for comments on the development of the project. This work was partially funded by the National Science Foundation award number CBET-2034495 RECODE and CBET-2145528 Faculty Early Career Development Program (L.M.); the National Institute of General Medicine of the NIH award number R35 GM138256 (L.M.);; grant 2023-332386 from the Chan Zuckerberg Initiative Donor Advised Fund, CZI DAF, an advised fund of the Silicon Valley Community Foundation (L.M., M.T.).

## Declaration of interests

L.M. is an inventor on a synNotch patent for applications in cancer cell therapy licensed to Gilead. The remaining authors declare no competing interests.

## Notes

https://github.com/TDEL-SyntheticBio/SyntheticElongation.git

