## Supplemental figures and figure legends for "Programming the elongation of mammalian cell aggregates with synthetic gene circuits"

### Supplementary Figure 1

**a** *In silico*

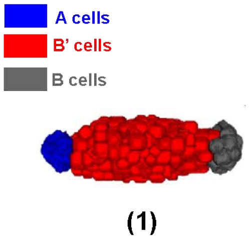


| | *Cell-cell adhesion strength* | | | ***Proliferation (daily growth rate)*** | **Cellular motility** | B→B' conversion rate |
|---|---|---|---|---|---|---|
| Cell type | A | B | B' | | | |
| A | strong | weak | weak | 0 | high | balanced |
| B | | strong | weak | 0.8 | high | |
| B' | | | strong | 0 | low | |

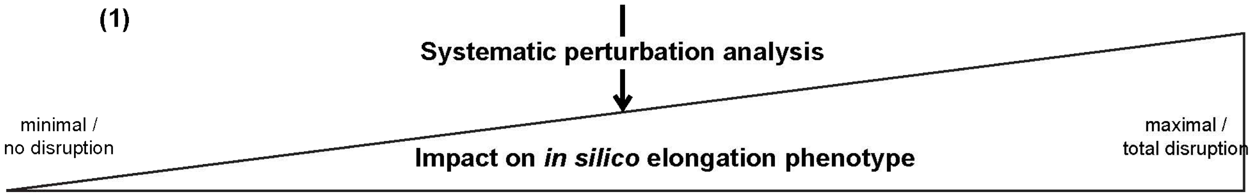


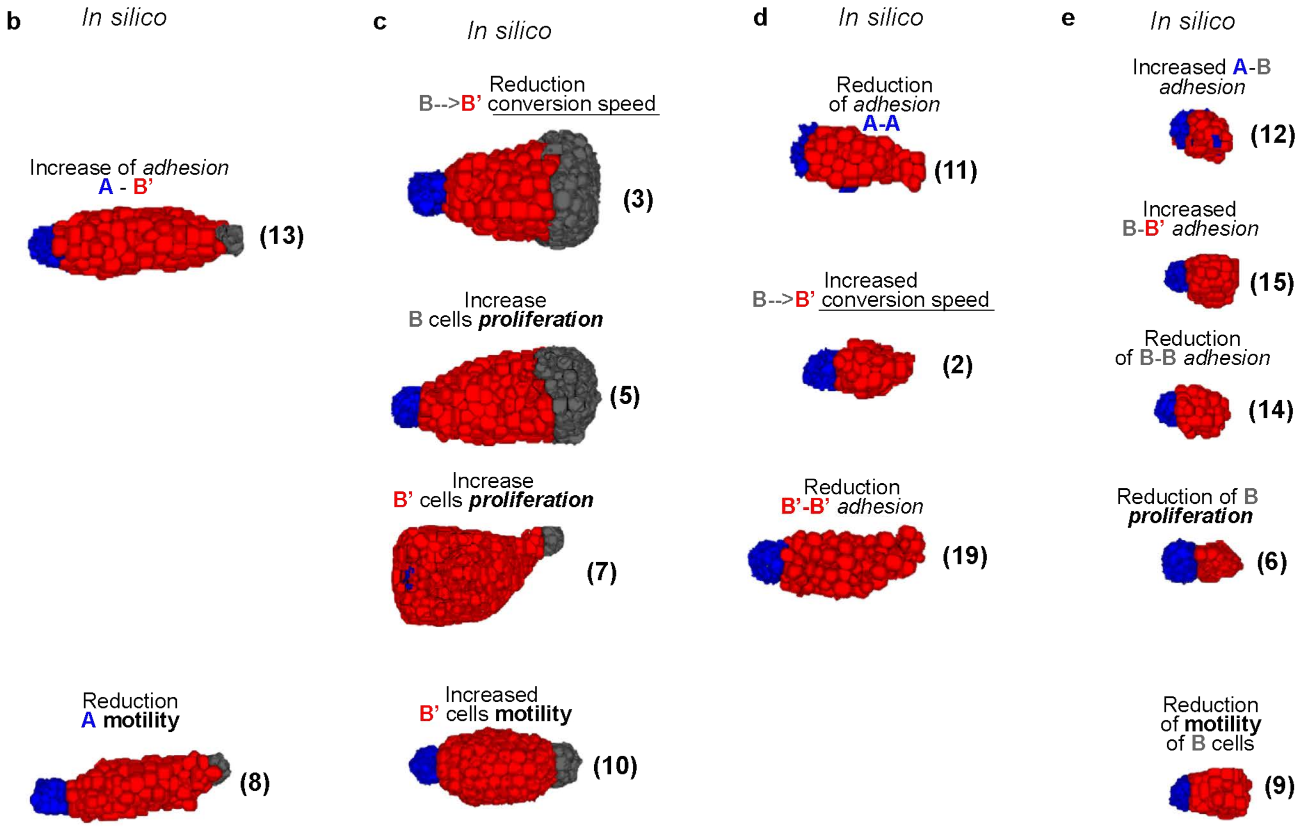


**f** *In silico*

| | *Cell-cell adhesion strength* | | |
|---|---|---|---|
| Cell type | A | B | B' |
| A | strong | weak | weak |
| B | | strong | weak/strong |
| B' | | | strong |

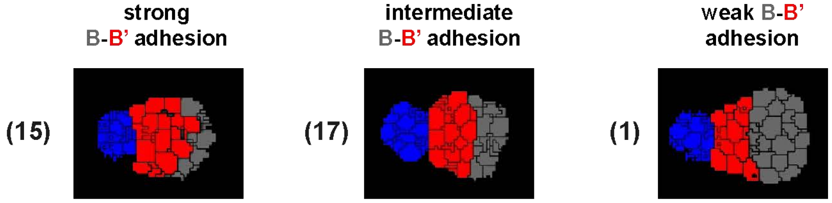

**Figure S1. *In silico* sensitivity analysis of elongation assay based on changes in selected parameters (related to Main Fig. 1).**

**a.** Left, 3D rendering of elongation assay *in silico* at day 7. Right, corresponding "genome", i.e. list of *in silico* parameters used for the simulation. For the corresponding numeric parameters used to set variables see Table. 1.1. In Table 1.1, the genomes are numbered as in this figure from (1) to (19). **b.** In vertical arrangement are two results of simulation of elongation assays specifically #13 and #8, which show relatively little perturbation of the elongation compared to the #1 implementation reference. **c.** Group of simulations generated with modification of parameters that generate substantial changes to the elongation phenotype. **d.** group of simulations generated with modification of parameters that generate elongation phenotypes but that have completely exhausted gray cells before the endpoint of simulation. **e.** group of simulations generated with modification of parameters that generate non-elongated phenotypes. Over each 3D rendering of the end-point elongation assay is written which parameter was modified and in which direction compared to the reference genome. For complete genome list see Table 1.1.

**f.** Left, qualitative adhesion matrix with parameters used to generate the simulations on the right. For complete numerical values see Table 1.1. Right, three examples of 2D sections of *in silico* simulations of elongation assays as described in the main text, for 3 different values of B-B'-type cells adhesion. The corresponding genomes are in Table 1.1, annotated with numbers as here.

#### Supplementary Figure 2

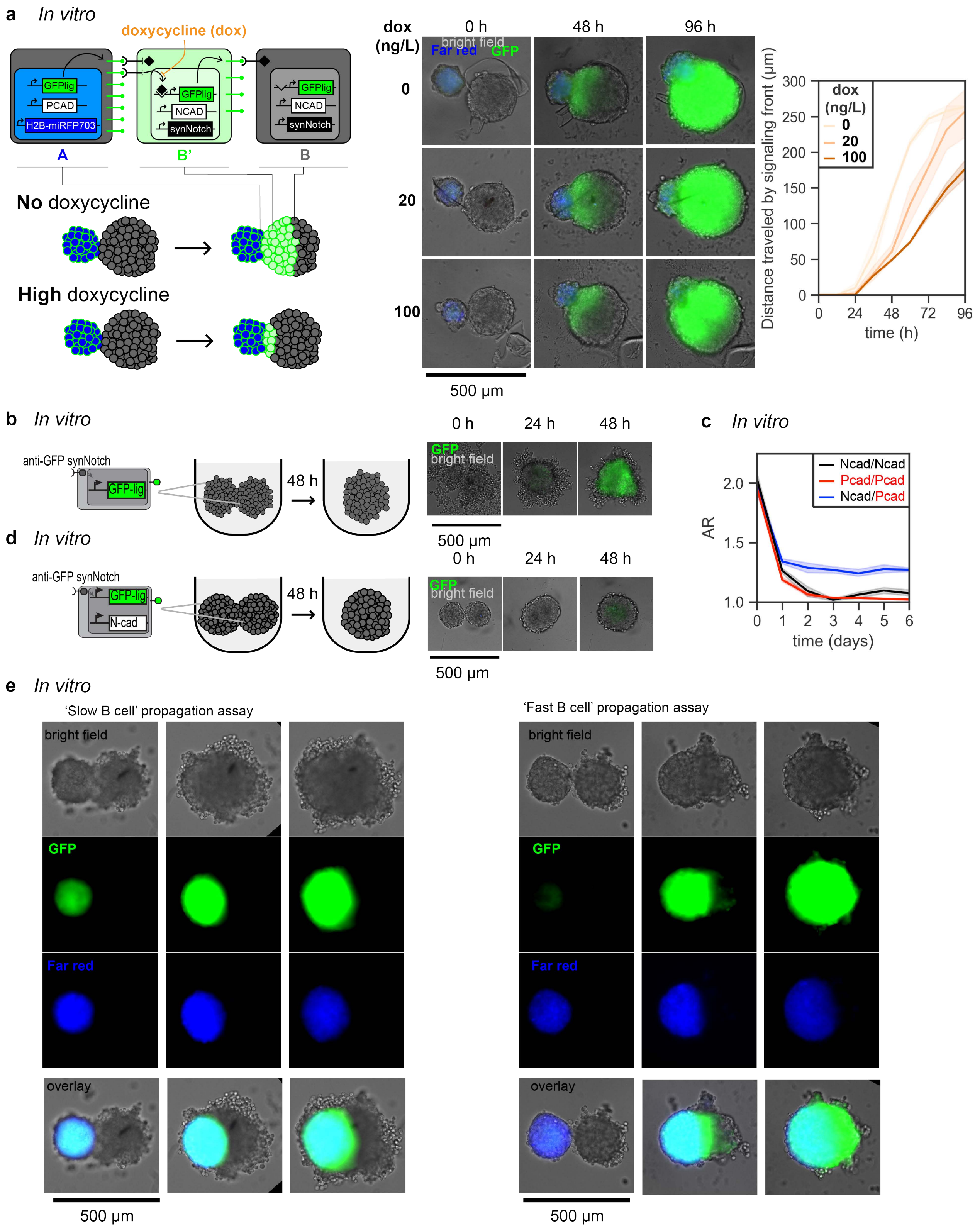

**Figure S2. Signal propagation assay and controls**

**a.** Signal propagation assay across spheroid of transceiver B-type cells *in vitro*. Left, schematic of the circuit, cells and genetic elements. Center, micrographs of representative *in vitro* assays. Far-red signal is rendered in blue (A-type cells), GFP signal is in green (B'-type cells), and brightfield is in gray (B-type cells do not express fluorescent markers). Scale bar is 500um. Right, graph of quantified signal propagation front over time in a signal propagation assay for the indicated conditions of Dox concentration.

**b.** Signal propagation assay – control for Fig. 4d-g. Left, schematic of cells, circuit and plating in wells. The two spheroids are both originated by transceiver B-type cells. Right, micrographs of representative *in vitro* assay. GFP signal is in green and inactivated B-type cells are in gray. Scale bar is 500um. **c.** Aspect ratio (AR) graph over time for the indicated spheroid co-cultures. Represents the same graph as in Main Fig. 4c but includes the day 0 value. **d.** Signal propagation assay – control for Main Fig. 4d-g. Left, schematic of cells, circuit, and plating in wells. Both spheroids are made by B-type cells that constitutively overexpress N-cadherin. Right, micrographs of representative *in vitro* assay. GFP signal is in green and inactivated B-type cells are in gray. Scale bar is 500um. See also videos SV7-8.

**e.** single-channel images of Main Fig. 3.

#### Supplementary Figure 3

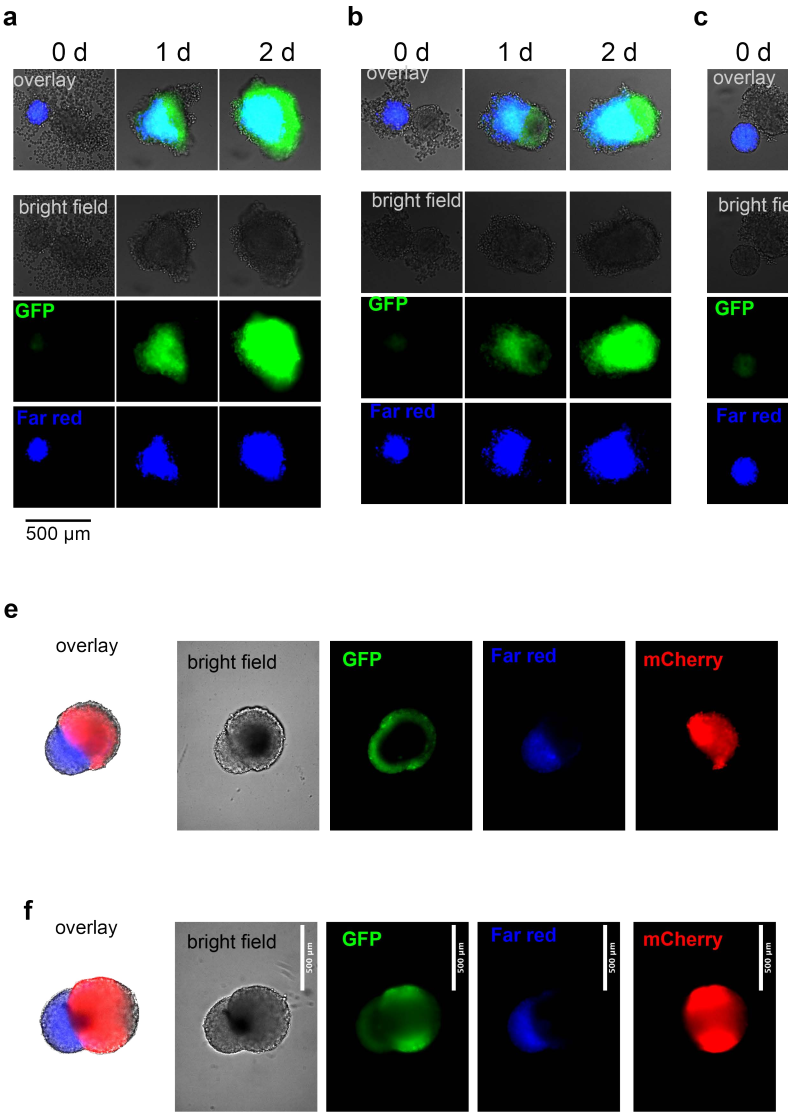

**Figure S3. Single channel images for Main Fig. 4**

**a.** Single channel images for Main Fig. 4d
**b.** Single channel images for Main Fig. 4e
**c.** Single channel images for Main Fig. 4f
**d.** Single channel images for Main Fig. 4g
**e.** Single channel images for Main Fig. 4i
**f.** Single channel images for Main Fig. 4k

#### Supplementary Figure 4

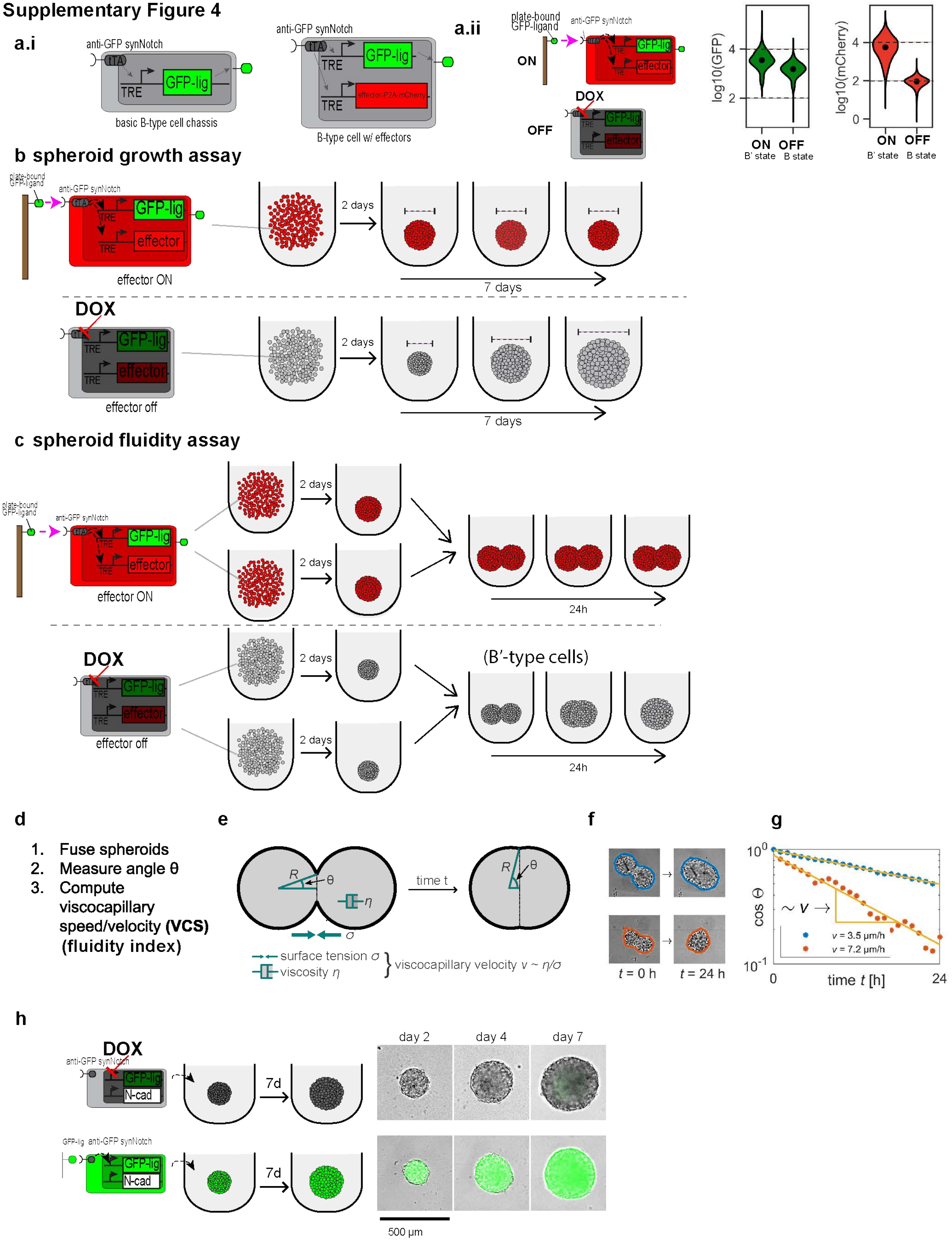

**Figure S4. Experimental pipeline for quantification of tissue growth and fluidity (related to main Fig. 5)**
**a.i.** Schematic of two B-type cells. Left, basic transceiver, that expresses anti-GFP-synNotch that drives GFP-ligand (green) expression. Right, transceiver with effectors: B-type cells express anti-GFP-synNotch driving GFP-lig, effector genes, and mCherry, separated from the effector in the same coding sequence by a P2A sequence (red).
**a.ii**: Left: schematics of generation of the ON (B'-type, top cell) and OFF (B-type, bottom cell) states. Right, FACS plots of cells: ON is after 3 days culture on GFP, OFF is after 3 days of culture in dox 10 mg/L. FACS data from ON/OFF transceivers, n=1 experiment, 4000-9000 cells/sample, log10 scale, point is median.
**b.** Schematic of tissue growth assay. Top row, condition of pre-activation. Brown line = scaffold surface; green ellipsoid = GFP ligand; pink arrow with dashed line = signaling interaction. Activated B'-type cells are shown in red. Measures of the diameter of the spheroids (dashed line) are taken at days 2, 4, and 7. Bottom row, signaling inhibition condition. Dox inhibits receptor activation, maintaining B-type cells in their basal B state; cells are collected in a U-bottom well, and similarly, the diameter of the resulting spheroid is measured every day for 7 days. Results of these assays with different cell lines are shown in Main Fig. 5a-c.
**c.** Schematic of spheroid fluidity assay. Top row, condition of pre-activation. Brown line = scaffold surface; green ellipsoid = GFP ligand; pink arrow with dashed line = signaling interaction. Activated B'-type cells are shown in red. 2 spheroids of B'-type cells are generated in 2 separate wells for 24h and then fused; at that point a 24-hour time-lapse is taken with images every 30' (see methods). Bottom row, signaling inhibition condition. Dox inhibits receptor activation, maintaining B-type cells in their basal B state; 2 spheroids of B-type cells are generated in 2 separate wells for 24-hour and then fused; at that point a 24-hour time-lapse is taken with images every 30'. Results of these assays with different cell lines are shown in Main Fig. 5d-f. **d-g.** Schematic with representative results of the pipeline for the measurement of the viscocapillary speed from the time-lapse imaging of fusing assembloids. **d.** List of steps. **e.** Schematic of fusion with measured quantities. **f.** Sample micrograph images for fusion assay of the B-type cell line with constitutive N-cad and no effectors (top, blue boundary, fuses slower), and B-type cell line with constitutive N-cad and p53 as an effector (orange boundary, fuses faster). **g.** Graph of cosTheta over time for the two samples from f., showing in yellow the fit used to measure the viscocapillary speed / velocity ('fluidity index').
**h.** Tissue growth assay – control for Main Fig. 5a-c. 1000 B-type cells are dispensed in a U-bottom well, and their growth is captured with imaging at days 2, 4, and 7. B/B'-type cells in this case constitutively express N-cad and are either prevented from signaling with dox (top row), or pre-activated before the growth assay (bottom row). On the right, representative microscope images of a spheroid of B-type cells in dox at the indicated time points. GFP signal is rendered in green, overlaid to the bright field. Scale bar is 500um.

Supplementary Figure 5

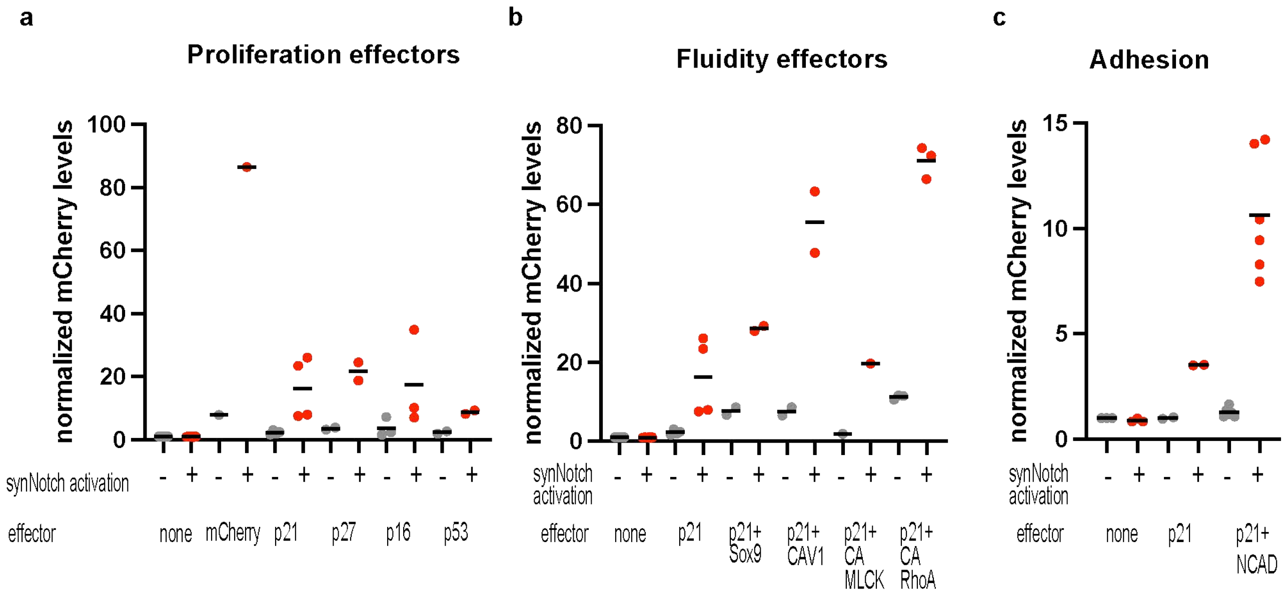

**Figure S5. mCherry expression levels in transceiver spheroids (related to main Fig. 5)**
**a.** Graph of average normalized mCherry levels from epifluorescence imaging of B/B'-type cells spheroids with the indicated effectors, 2 days following seeding after either stimulation with plate-bound GFP ligand (B') or culture in doxycycline (B). B/B' cells are transceiver cells with constitutive N-cad, as the ones depicted schematically in Main Fig. 5a-b. Experimental averages from n=1-4 individual experiments. Each experimental average was generated from at least 3 individual spheroids. For mCherry normalization procedure see Methods.
**b.** Graph of average normalized mCherry levels from epifluorescence imaging of B/B'-type cells spheroids with the indicated effectors, 2 days following seeding after either stimulation with plate-bound GFP ligand (B') or culture in doxycycline (B). B/B'-type cells are transceiver cells with constitutive N-cad as the ones depicted schematically in Main Fig. 5d. Experimental averages from n=1-4 individual experiments. Each experimental average was generated from at least 3 individual spheroids. For mCherry normalization procedure see Methods.
**c.** Graph of average normalized mCherry levels from epifluorescence imaging of B/B'-type cells spheroids with the indicated effectors, 2 days following seeding after either stimulation with plate-bound GFP ligand (B') or culture in doxycycline (B). B/B'-type cells are transceiver cells with no constitutive adhesion protein expressed, as the ones depicted schematically in Main Fig. 2b. Experimental averages from n=2-6 individual experiments. Each experimental average was generated from at least 3 individual spheroids. For mCherry normalization procedure see Methods.
For all experiments, to generate B' state cells, cells were stimulated with plate-bound GFP as described in the methods for 24h, then a spheroid made from 1000 cells was formed for 24h and then image analysis is performed. A control condition per each cell line is generated with treatment of dox and no exposure to GFP ligands (B-type cells). Gray dots are for the un-stimulated conditions, red dots are for the conditions with stimulation. Short black horizontal lines denote the average of multiple experiments.

**Supplementary Figure 6**

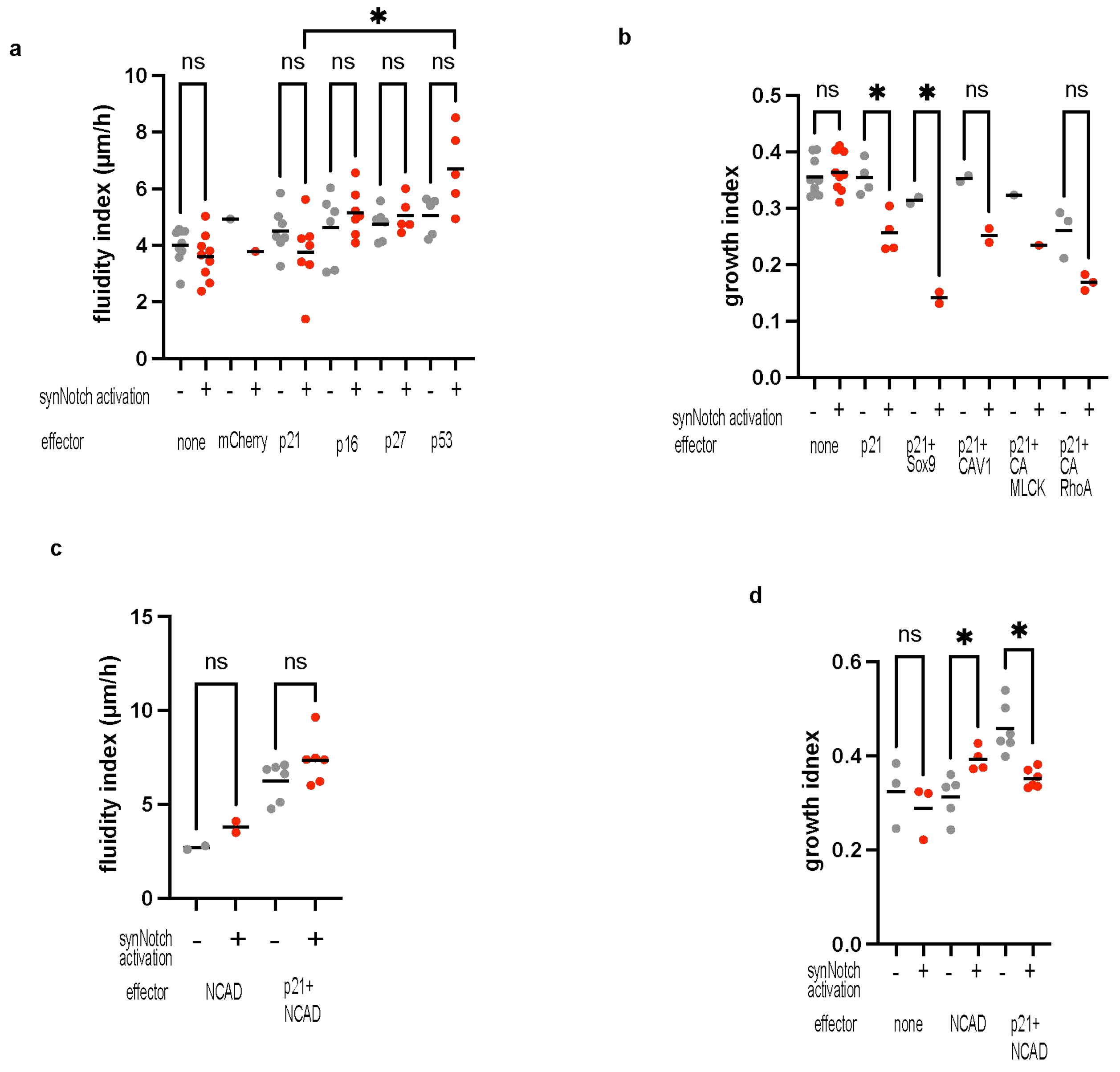

**Figure S6. Impact of growth effectors on tissue fluidity and of fluidity effectors on tissue growth (related to main Fig. 5)**

**a.** Graph of fluidity index of B/B'-type cells spheroids with the indicated proliferation effectors. Fluidity index is calculated with the spheroid fluidity assay described in Fig. S3 and reported also in Main Fig. 5d-f. The B/B'-type cells analyzed here are transceiver cells with constitutive N-cad, as the ones depicted schematically in Main Fig. 5a-b. Experimental averages from n=1-7 individual experiments, with each experimental average generated from at least n=2 technical replicates. **b.** Graph of the daily growth rate of B-type cells spheroids with the indicated fluidity and proliferation effectors. Daily growth rate index is calculated with the spheroid growth assay as described in Fig. S3 and reported also in Main Fig. 5a-c. The B/B'-type cells analyzed here are transceiver cells with constitutive N-cad and with fluidity effectors, as the ones depicted schematically in Main Fig. 5d. Experimental averages from n=1-7 individual experiments, with each experimental average generated from at least n=2 technical replicates.

**c.** Graph of fluidity index of B/B'-type cells spheroids with the indicated adhesion and proliferation effectors. Fluidity index is calculated with the spheroid fluidity assay described in Fig. S3 and reported also in Main Fig. 5d-f. The B/B'-type cells analyzed here are transceiver cells without any constitutive adhesion protein, as the ones depicted schematically in Main Fig. 2b. Experimental averages from n=1-6 individual experiments, with each experimental average generated from at least n=2 technical replicates.

**d.** Graph of the daily growth rate of B-type cells spheroids with the indicated adhesion and proliferation effectors. Daily growth rate index is calculated with the spheroid growth assay as described in Fig. S3 and reported also in Main Fig. 5a-c. The B/B'-type cells analyzed here are transceiver cells without any constitutive adhesion protein, as the ones depicted schematically in Main Fig. 2b. Experimental averages from n=1-6 individual experiments, with each experimental average generated from at least n=2 technical replicates.

For **a, b, c,** and **d.** Statistical comparisons were performed with Brown-Forsythe and Welch ANOVA tests with selected multiple comparisons, * indicate $p<0.1$ and ns nonsignificant comparisons.

**Supplementary Figure 7**

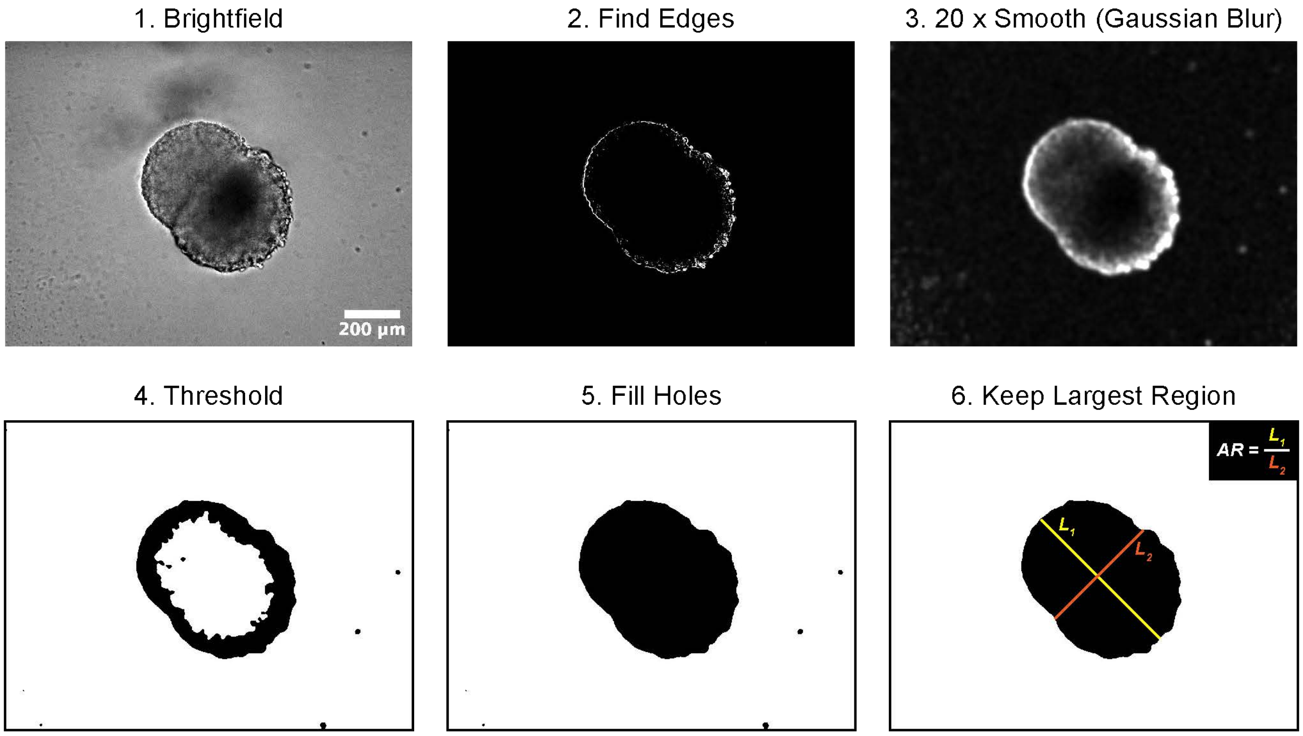

**Figure S7. Pipeline for automated quantification of tissue elongation.**
Pipeline for image analysis for quantification of the aspect ratio of spheroids and assembloids. The numbered items indicate Fiji functions used at each step, see methods for details. Aspect ratio (AR) is calculated as the ratio between the longest axis (L1, yellow line) and the shortest axis (L2, orange line) of the inbounding ellipsoid.

**Supplementary Figure 8**

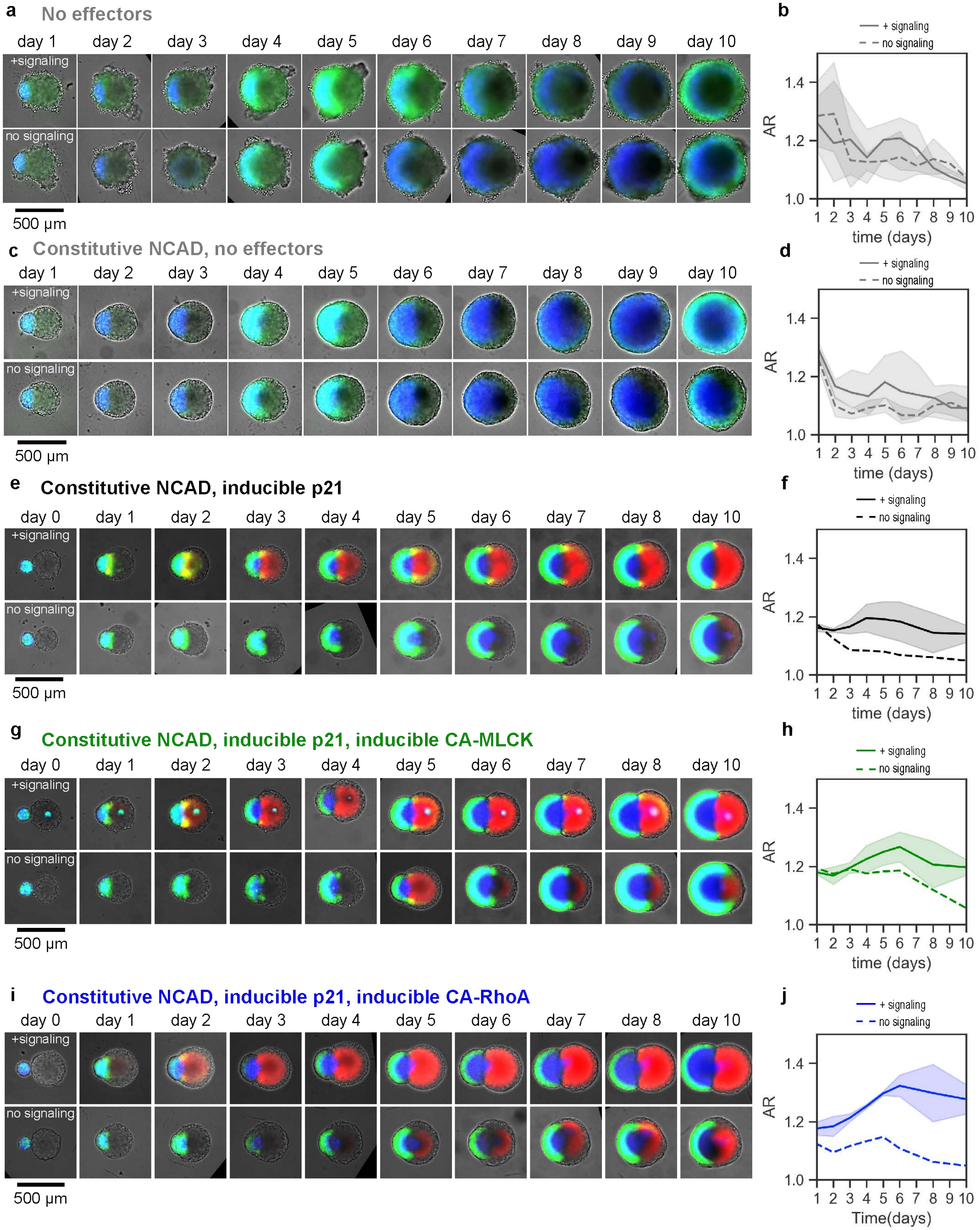

**Figure S8. Time series of elongation experiments – related to Main Fig. 6**

**a, c, e, g, i.** Daily micrographs from day 1 to day 10, of the dataset used in the Figure 6 illustrations; constructs are generated following the fusion of a 200 cells spheroid of Pcad+ sender cells with a 4000 cells spheroid of transceivers as follows:
**a**: transceivers with no effectors downstream of synNotch activation (shown also in Fig. 6b)
**c:** transceivers with constitutive Ncad expression and no effectors downstream of synNotch activation
**e**: transceivers with constitutive Ncad expression and p21 induction upon synNotch activation (shown also in Fig. 6d)
**g**: transceivers with constitutive Ncad expression and p21 + CA-MLCK induction upon synNotch activation (shown also in Fig. 6f)
**i**: transceivers with constitutive Ncad expression and p21 + CA-RhoA induction upon synNotch activation. (shown also in Fig. 6h)
Green: GFPlig (senders, activated transceivers); blue: miRFP703 (senders); red: mCherry (activated transceivers with effectors). +signaling condition: transceivers were culture with a low amount of tetracycline up to day 0, when they were fused with senders. No signaling condition: transceivers were cultured similarly to the +signaling condition up to day 0, after which the spheroids were kept in 10 mg/L doxycycline to inhibit synNotch signaling.

**b, d, f, h, j.** AR quantifications corresponding to the conditions in a, c, e, g, i, respectively. **b** and **d.** n=1 experiment (n=3-6 technical replicates). Error bars: s.d. **f, h and j.** n=1 experiment for the no signaling condition, n=2 experiments for the +signaling condition. Experimental averages were obtained from a minimum of 3 technical replicates. Error bars: s.e.m.

**Supplementary Figure 9**

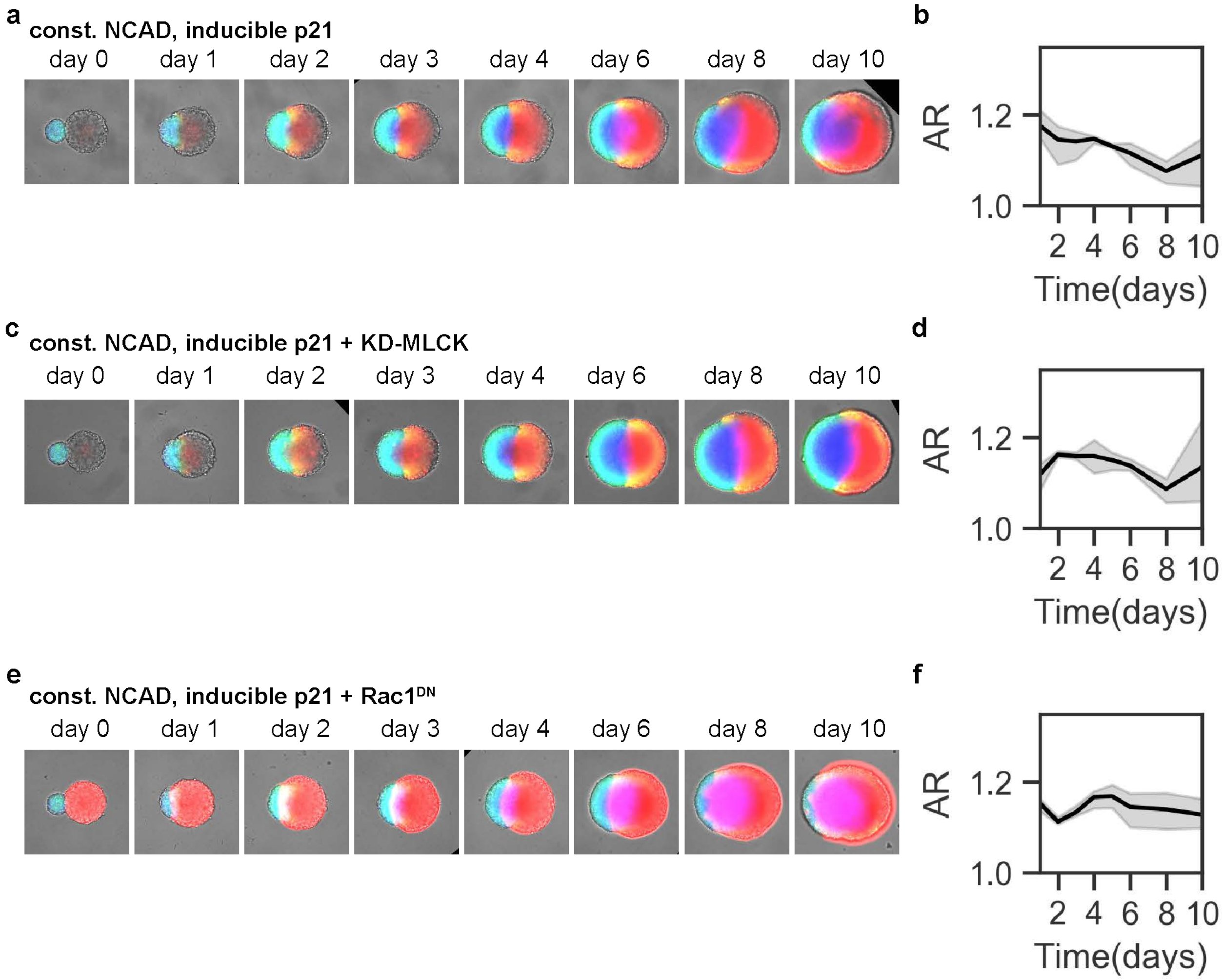

**Figure S9. Time lapse of elongation experiments that do not display an increase in the AR index over time (related to main Fig. 6)**

**a, c, and e.** Daily micrographs of a single experiment following the fusion of a 200-cell spheroid of P-cad+ sender cells with a 4000-cell spheroid of the following transceiver types:
**a**: transceivers with constitutive Ncad expression and p21 induction upon synNotch activation,
**c**: transceivers with constitutive Ncad expression and p21 + KD-MLCK induction upon synNotch activation,
**e**: transceivers with constitutive Ncad expression and p21 + $Rac1^{DN}$ induction upon synNotch activation.
Green: GFPlig (senders, activated transceivers); blue: miRFP703 (senders); red: mCherry (activated transceivers with effectors). All data provided are generated in the +signaling condition: transceivers were culture with a low amount of tetracycline up to day 0 when they are fused with senders.

**b, d, and f.** AR quantifications correspond to the conditions in a, c and e, respectively. n=1 experiment, n=3 technical replicates. Error bars: s.d

#### Supplementary Figure 10

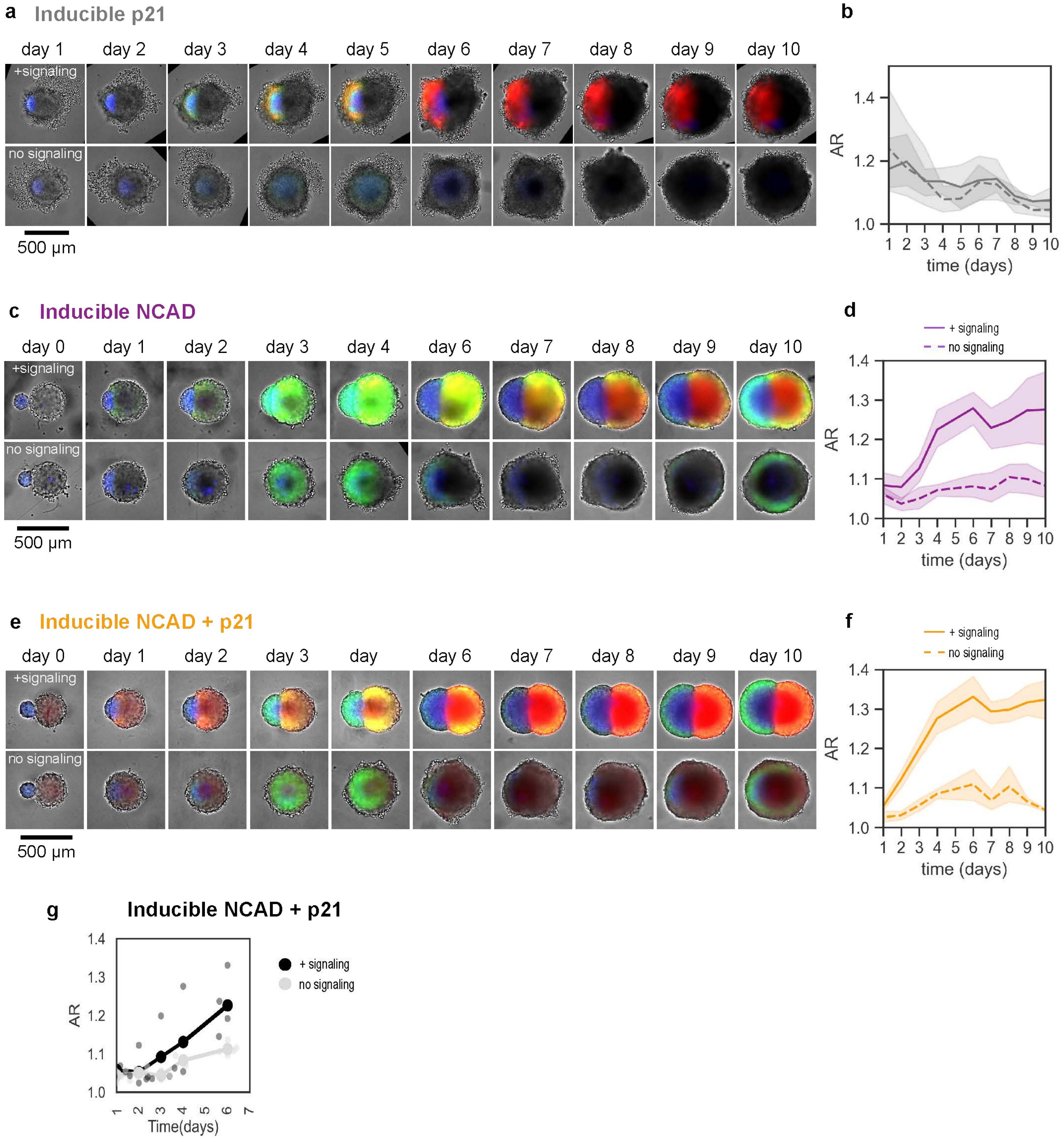

**Figure S10. Time series of elongation experiments – related to Main Fig. 7**

**a, c and e.** Daily micrographs of a single experiment following the fusion of a 200-cell spheroid of P-cad+ sender cells with a 4000 cells spheroid of transceiver cells of the following types:
**a**: transceivers with p21 induction upon synNotch activation,
**c**: transceivers with Ncad induction upon synNotch activation (also shown in Main Fig. 7d),
**e**: transceivers with Ncad + p21 induction upon synNotch activation (also shown in Main Fig. 7f).
Green: GFPlig (senders, activated transceivers); blue: miRFP703 (senders); red: mCherry (activated transceivers with effectors). +signaling condition: transceivers were culture with a low amount of tetracycline up to day 0, when they are fused with senders. No signaling condition: transceivers were cultured similarly to the +signaling condition up to day 0, after which the spheroids were kept in 10 mg/L doxycycline to inhibit synNotch signaling.

**b, d and f.** AR quantifications correspond to the conditions in a, c, and e, respectively. n=1 experiment, n=3-6 technical replicates. Error bars: s.d

**g.** Aspect ratio graph over time for structures generated with 200 cells spheroid of Pcad+ sender cells with a 4000 cells spheroid of transceiver with Ncad + p21 induction upon synNotch activation. Shows experimental averages as individual data points, n=4 experiments.

#### Supplementary Figure 11

a

Tissue growth can be controlled through mitosis rate (growth variable) *in silico*

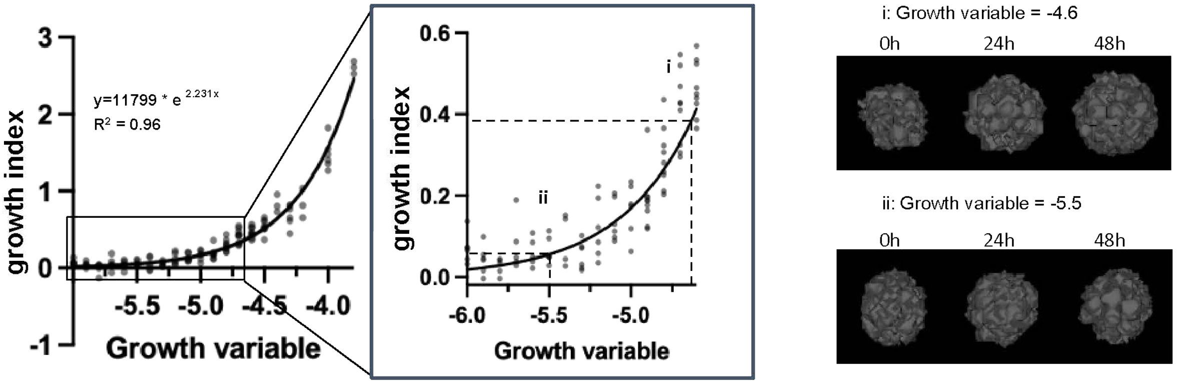


b Control over tissue rounding speed *in silico*

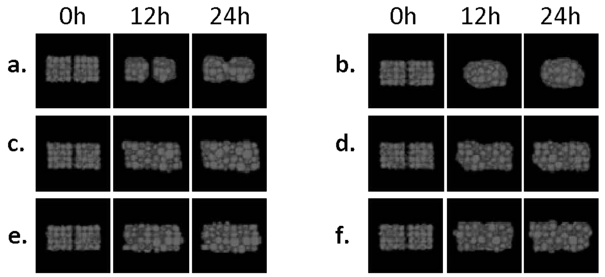


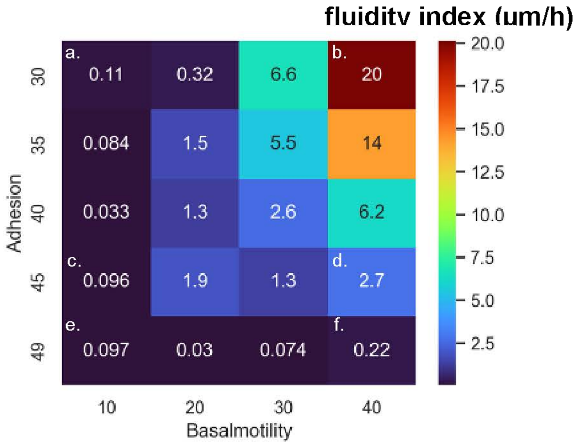


c impact of proliferation on tissue fluidity *in silico*

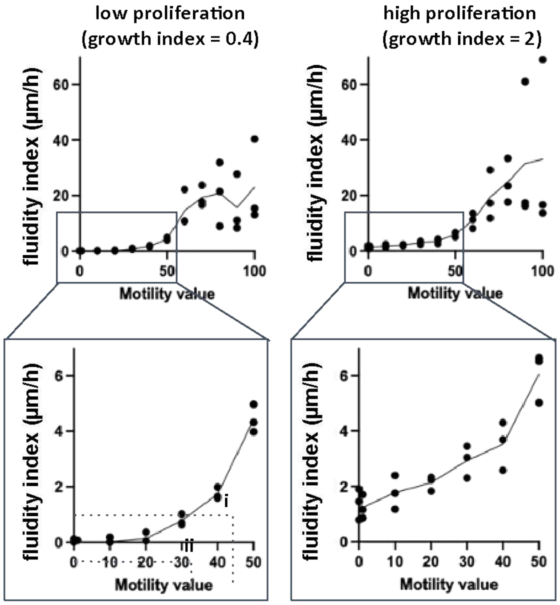


d impact of motility on tissue growth *in silico*

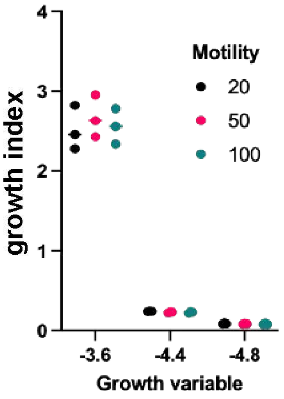

**Figure S11. Parametrization of the computational model**

**a.** Left, graph of daily growth rate over a range of growth variable values; black rectangle is enlarged in the inset. In the inset, i and ii mark specific instances of growth variable values that are shown on the right part of the panel. Right, time course of 48h of simulated time of spheroid growth. The two simulations are run keeping all the other parameters equal, varying only the growth variable as indicated. Growth index is calculated in the same way as for the *in vitro* spheroids, as depicted in Fig. S3 and as described in the methods, from measures of the diameter of the spheroid over time. **b.** Left, depiction of timepoints of simulated homotypic spheroid fusion assays *in silico*. Two spheroids of 64 cells approximately are juxtaposed, and their evolution is followed over 24h of simulated time. The images are then taken every 30’ and then processed in the same way as for the in vitro microscope images, as described in Fig. S3d. The 6 conditions a.-f. have parameters as indicated in the matrix on the right. Right, fluidity index (viscocapillary speed, um/h), measured from the spheroid fusion assay in silico across different parameters of adhesion and of basal motility as indicated. On the right is the color-code corresponding to the different levels of fluidity index; individual values are also written in the respective matrix cells. The average of n=6 simulations is shown for each condition. **c.** Graphs of fluidity index values from spheroid fusion assays *in silico* over a range of basal motility values in two conditions: low proliferation at (left) or high proliferation (right) with the indicated growth index. For both graphs, the black rectangle is enlarged in the bottom inset. **d.** Graphs of growth index for spheroids generated with cells varying across 3 levels of basal motility, and 3 levels of proliferation rate.

#### Supplementary Figure 12

**a**

| **gene circuit ID** | ***morphospace axis***<br>**Phase separation between B and B' (estimate)** | **Proliferation change after B→B' conversion (DGR^B/DGR^B')** | **Resistance to rounding of B' (VCS^B')** | *time of in vitro snapshot shown in figure (hours)* |
|---|---|---|---|---|
| ***in silico realizations for morphospace extremes*** | | | | |
| **000** | none | 1 | 0.11 | |
| **001** | none | 1 | 1 | |
| **010** | none | 12 | 0.11 | |
| **100** | high | 1 | 0.03 | |
| **011** | none | 12 | 1 | |
| **101** | high | 1 | 1 | |
| **110** | high | 12 | 0.03 | |
| **111** | high | 12 | 1 | |
| ***in silico simulations corresponding to specific in vitro realizations*** | | | | |
| **a: const. NCAD** | none | 1 | 0.4 | 101 |
| **b: induc. p21** | none | 1.4 | 0.03 | 85 |
| **c: induc. p21 + NCAD** | intermediate | 1.4 | 0.14 | 131 |
| **d: const. NCAD, induc. p21** | none | 1.3 | 0.29 | 95 |
| **e: const. NCAD, induc. p21 + CA-MLCK** | none | 1.32 | 0.36 | 164 |
| **f: const. NCAD, induc. p21 + CA-RhoA** | none | 1.59 | 1 | 151 |

**b**

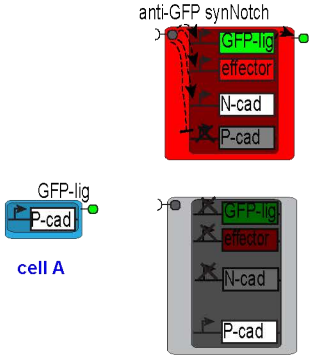


**c**

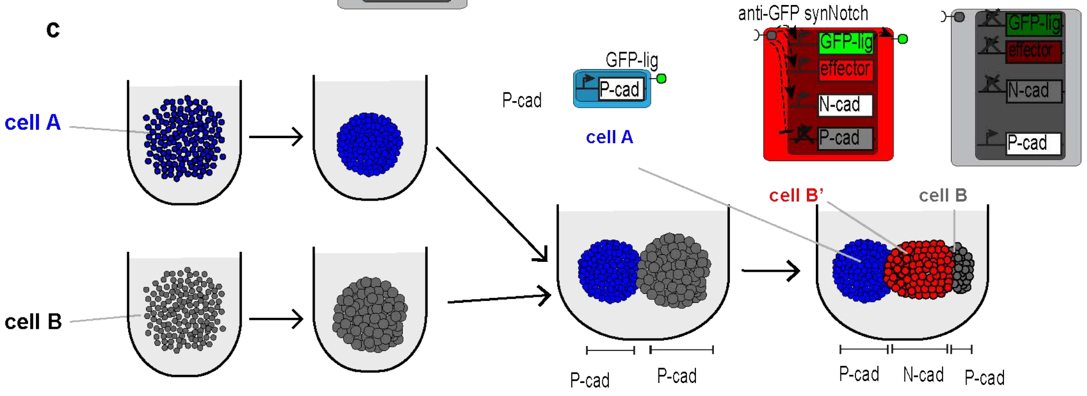

**Figure S12. Morphospace additional information**

**a.** Coordinates along the 3 morphospace axes for the *in silico* simulations shown in Main Fig. 8. For the morphospace extremes list, the id of genomes here corresponds to numbers shown in white on top of simulated 3D structures in Main Fig. 8. For a complete list of all genomes with quantitative parameters see Table 1.2. For the *in silico* genomes corresponding to specific in vitro realizations, the id of genomes here corresponds to letters shown in white on top of simulated 3D structures in Main Fig. 8. For a complete list of all genomes with quantitative parameters see Table 1.3. **b.** Schematic depiction of the predicted circuit for stronger elongation *in vitro*. **c.** Schematic of expected behavior *in vitro* of the genetic circuit shown in b.
